# Re-emergence and diversification of a specialised antennal lobe morphology in ithomiine butterflies

**DOI:** 10.1101/2020.10.13.336206

**Authors:** Billy J Morris, Antoine Couto, Asli Aydin, Stephen H Montgomery

## Abstract

How an organism’s sensory system functions is central to how it navigates its environment and meets the behavioural challenges associated with survival and reproduction. Comparing sensory systems across species can reveal how facets of behaviour and ecology promote adaptive shifts in the relative importance of certain environmental cues. The insect olfactory system is prominent model for investigating how ecological factors impact sensory reception and processing. Notably work in Lepidoptera led to the discovery of vastly expanded structures, termed a macroglomerular complex (MGC), within the primary olfactory processing centre. These structures typically process pheromonal cues and provide a classic example of how variation in size can influence the functional processing of sensory cues. Though prevalent across moths, the MGC was lost during the early evolution of butterflies, consistent with evidence that courtship initiation in butterflies is primarily reliant upon visual cues, rather than long distance olfactory signals like pheromones. However, a MGC has recently been reported to be present in a species of ithomiine, *Godryis zavaleta*, suggesting this once lost neural adaptation has re-emerged in this clade. Here, we show that MGC’s, or MGC-like morphologies, are indeed widely distributed across the ithomiine tribe, and vary in both structure and the prevalence of sexual dimorphism. Based on patterns of variation across species with different chemical ecologies, we suggest that this structure is involved in the processing of both plant and pheromonal cues, of interlinked chemical constitution, and has evolved in conjunction with the increased importance and diversification of plant derived chemicals cues in ithomiines.

## Introduction

An organism’s sensory system is its interface with the rest of the world, the link between its internal and external environments. The manner in which sensory systems vary can reveal how different species are attuned to different cues, the association between cues and behaviour, and how behavioural variation maps to the evolution of sensory systems. Lepidopterans have often been used as models to investigate how ecological variability affects the evolution of olfactory systems (Carlsson et al. 2011; Bisch-Knaden et al. 2012; Carlsson et al. 2013; Namiki et al. 2014; van Dijk et al. 2017), and how the central brain processes olfactory information (Kanzaki et al. 1989; Anton and Hansson, 1994; Tabuchi et al. 2013). This includes classic work characterising pheromones, the olfactory response to these chemical cues, and the manner in which the pheromone processing system evolves (Butenandt et al. 1959; Klun and Maini, 1979; Namiki et al. 2014). Studies on lepidopteran sensory systems have provided crucial insights into how sensory systems function, how separate strands of information are processed and integrated within the brain, and the relationship between sensory systems and ecological variables (Couto et al. 2020).

As in all insects, the primary olfactory processing structure within the lepidopteran brain is the antennal lobe (AL). The AL is composed of a collection of functional and morphological units, termed glomeruli. Each glomerulus is a synapse dense region composed of the axon terminals of antennal sensory neurons that typically express the same olfactory receptor (Vosshall et al. 2000), local interneurons that refine the olfactory message, and projection neurons that convey information to higher brain centres. Odorants elicit activity across a range of olfactory receptors, and associated glomeruli, encoding the odorant identity through the combinatorial activation of glomeruli (Joerges et al. 1997; Galizia et al. 1999; Carlsson et al. 2002; Wang et al. 2003; Hallem and Carlson, 2006; Zube et al. 2007). Despite their ecological diversity, within Lepidoptera the antennal lobe is relatively consistent in its structure, being made up of ~60-80 glomeruli (Rospars, 1983; Berg et al. 2002; Kazawa et al. 2009; Heinze and Reppert, 2012; Montgomery and Ott, 2015; Montgomery et al. 2016; Zhao et al. 2016). However, in moths a prominent morphologically distinct sub-cluster of glomeruli occur at the base of the antennal nerve (Bretschneider, 1924; Matsumoto and Hildebrand, 1981; Koontz and Schneider, 1987). This glomerular cluster is termed a Macroglomerlar complex (MGC) and is composed of enlarged, ‘macro’ glomeruli (MG), and smaller, associated glomeruli, termed ‘satellite’ glomeruli. These glomeruli often display an extreme degree of sexual dimorphism, being vastly enlarged in males relative to females (Matsumoto and Hildebrand, 1981; Koontz and Schneider, 1987). Though first identified in moths, MGs/MGCs have subsequently been observed in a diverse range of insects, including in Blattodea, Diptera, Hymenoptera, and Lepidoptera (Chambille et al. 1980; Couto et al. 2016; Kelber et al. 2009; Ibba et al. 2010; Keubler et al. 2010). MGCs are typically involved in processing pheromonal cues detected by the antennal sensilla, where their corresponding olfactory receptors are highly expressed in a greater number of sensory neurons, providing heightened sensitivity (Warner et al. 2007; Miura et al. 2009). MGs responsive to host plant related cues have also been reported (Ibba et al. 2010), suggesting they reflect an efficient way of increasing sensitivity to biologically important odours to each species. MGs are therefore classic examples of how neuropil size reflects functional performance, as variation in their volume is generally associated with variation in sensitivity to their corresponding odour (Gronenberg and Hölldobler, 1999; El Jundi et al. 2009b; Warner et al. 2007; Miura et al. 2009). Furthermore, MGC structure and composition is variable across closely related species, suggesting they may co-evolve adaptively with species specific ecological and behavioural traits (Kondoh et al. 2003; Namiki et al. 2014; Bastin et al. 2018).

While MGCs are ubiquitous in moths (Rospars and Hildebrand, 2000; Huetteroth and Schachtner, 2005; El Jundi et al., 2009b; Løfaldli et al., 2010; Yan et al. 2019), including diurnal species (Stöckl et al. 2016), they are absent in several phylogenetically disparate butterflies (Rospars, 1983; Carlson et al. 2011; Heinze and Reppert, 2012; Montgomery et al. 2016) suggesting they were lost at the origin of Papilionoidea. This has been interpreted as reflecting an increased reliance on visual cues, and the decreased importance of long-distance chemical signalling in butterfly mating behaviours (Rospars, 1983; Rutowski, 1991; Andersson et al. 2007). However, this view is being revisited. Evidence is accumulating that pheromone cues function in interspecific discrimination, sexual attraction and discrimination, and expediate female acceptance in courtship in a range of butterflies (Andersson et al. 2007; Constanzo and Monteiro, 2007; Schulz et al. 2007; Mérot et al. 2015; Chengzhe et al. 2017; Darragh et al. 2017).

One diverse tribe of diurnal butterflies with particular reliance on olfactory cues are the Ithomiini. Ithomiines utilise derivatives of pyrrolizidine alkaloids (PAs) for both chemical defence and intraspecific communication (Pliske, 1975, Pliske et al. 1976; Brown, 1984). PAs are sequestered from particular species of plants at the adult stage, with males being significantly more attracted and motivated by these resources (Pliske, 1975; Brown 1984). Males provision eggs with PAs through the spermatophore, providing chemical protection to the egg and larvae (Brown 1984). PA derived pheromones are secreted from ‘hair pencils’, specialised, elongated cells found on the dorsal surface of the androconial gland on the forewing (Schulz et al.1988). The expression of PA-derived pheromones is believed to represent an honest signal of male quality and facilitates mating receptivity in females (Boppré, 1978). Male pheromones have been shown to serve a variety of functions across ithomines, acting purely as an attractant in some species, and as both a short-range female-attractant and long-range male-repellent in others (Pliske, 1975). Consistent with these differences, pheromone blends vary qualitatively and quantitatively across the tribe (Edgar et al. 1976; Brown, 1984; Brown 1987; Schulz 1988; Trigo et al. 1994; Trigo et al. 1996; Schulz et al. 2004; Stamm et al. 2019), and male mating strategies range from establishing and defending territories, to gregarious leks, or aggressive ‘take downs’ of females (Pliske, 1975).

The strong sexual dimorphism in adult attraction to PA sources, and the utilisation of these chemicals for both chemical defence and pheromones, suggests that olfactory adaptations to detect these compounds have had particular importance in ithomiine evolution. Indeed, the first sexually dimorphic MGC recorded in butterflies was recently described in an ithomine, *Godyris zavaleta* (Montgomery and Ott, 2015). This observation was surprising given evidence MGCs were lost early in butterfly evolution, with even species within the relatively closely related Danainae subfamily lacking MGCs (Heinze and Reppert, 2012). This strongly suggests the independent, convergent evolution of a once lost neural adaptation in this tribe, a striking example of a reversal in a phylogenetic trend. However, it remains unclear how prevalent MGCs are across ithomiines, whether this structure functions solely for the location of PA sources, or whether it is used for long-distance pheromone detection (Montgomery and Ott, 2015). Here, we utilise a comparative approach, sampling a further twelve, diverse species of ithomiine, to investigate these questions.

## Methods

### i) Animals

Specimens were collected in the Estación Científica Yasuní, in the Parque Nacional Yasuní, Orellana Province, Ecuador, during November-December 2011, and September-October 2012, under permit 0033-FAU-MAE-DPO-PNY and exported under permits 001-FAU-MAE-DPO-PNY and 006-EXP-CIEN-FAU-DPO-PNY. Permits were obtained from Parque Nacional Yasuní, Ministerio Del Ambiente, La Dirección Provincial de Orellana. Species representing 12 genera, excluding *Godyris*, were selected on the basis of phylogenetic distribution and available sample size, and represent 8 of the 10 ithomiini subtribes **(Table S1)**. Dissection and fixation of specimens were performed at the Estación Científica Yasuní. Brains were exposed by removing a section of head carapace under HEPES-buffered saline (HBS; 150 mM NaCL; 5mM KCL; 5 mM CaCl_2_; 25 mM sucrose; 10mM HEPES; ph 7.4), before being fixed with zinc formaldehyde solution (ZnFA; 0.25% [18.4 mM] ZnCl_2_; 0.788% [135mM] NaCl; 1.2% [35mM] sucrose; 1% formaldehyde) for 16-20 hours. Extraneous head tissue was then removed, and brains were washed three times in HBS. Samples were transferred to 80% methanol/20% DMSO for at least two hours, then stored in 100% methanol. Samples were kept at room temperature until return to the UK, then transferred to −20°C.

### ii) Immunohistochemistry

Samples were rehydrated using serial Tris buffer-methanol solutions (90%, 70%, 50%, 30% and 0%), each for 10 minutes. Brains were then incubated for two hours in NGS-PBS_d_ (5% Normal Goat Serum, 1% DMSO, 94% 0.1M PBS), before being exposed to anti-SYNORF1 (Antibody 3C11; Developmental Studies Hybridoma Bank, University of Iowa, Iowa City, IA, RRID: AB_2315424; Buchner, 1996) in solution with PBS_d_-NGS, at a ratio of [1:30], for 3.5 days at 4°C. Non-bound antibody was removed by washing with PBS_d_ (1% DMSO, 99% 0.1M PBS) three times. Goat anti-mouse secondary antibody, Cy2-conjugated (Jackson ImmunoResearch; Cat No. 115–225-146, RRID: AB_2307343, West Grove, PA), was then applied at [1:100] in PBS_d_-NGS for 2.5 days at 4°C. Samples were imbued with glycerol through graded exposure in 0.1M Tris buffer (1%, 2%, 4% each for two hours, and 8%, 15%, 30%, 60%, 70%, 80%, each for 1 hour), and full dehydrated by washing with 100% ethanol (3 x 30 minutes). Methyl Salicylate was then underlaid, and the brains allowed to sink. This was repeated twice before transfer to storage vials of methyl salicylate.

### iii) Confocal imaging and Image Segmentation

Samples were mounted in methyl salicylate between two cover slips, either side of a hole bored through an aluminium slide. *Mechanitis* and *Ithomina* were imaged on a Leica SP5 microscope using a 10x 0.4NA objective. All other species were scanned on an Olympus IX3-SSU using a 10x 0.4NA objective. As we do not compare raw volumes across species, the use of different microscopes does not affect our subsequent analyses. For each individual, a single stack was taken encompassing the whole AL with a z-step of 1μm between each optical section and a *x-y* resolution of 1024×1024 pixels. Consistent light detection was ensured by adjusting the laser intensity and gain with depth. To correct for the artefactual shortening of the z-dimension of images due to the air objective lenses, a correction factor of 1.52 was applied to the *z*-dimension of the image stacks (Heinze and Reppert, 2012; Montgomery and Ott, 2015). Image segmentation was performed in Amira 5.4.1 (ThermoFisher Scientific; RID: SCR_007353). Volumes of evaluated areas were exported using the *measure statistics* tool. Volumes for surface models were plotted using the *nat* R package (Bates et al. 2020). Putative MGCs were identified on the basis of internal fibrous structure and location at the base of the antennal nerve, as described in Montgomery and Ott (2015). To formally examine the presence of MGs, each individual glomerulus was segmented in two focal male ALs for each species. In *Methona* 4 males were analysed in this way to confirm the apparent lack of both MGs and MGCs (see results). Males were chosen for this initial assessment as the MGs are larger in *Godyris* males (Montgomery and Ott, 2015), as is commonly observed across moths (Rospars and Hildebrand, 1992; Huetteroth and Schachtner, 2005; El Jundi et al., 2009b; Løfaldli et al., 2010; Yan et al. 2019). Subsequent visual inspection of females did not reveal any glomeruli specifically enlarged in females. Three tests were used to evaluate whether MGs were present in focal individuals, based on methodologies used in previous publications (Montgomery and Ott, 2015; Keubler et al, 2010; Kelber et al. 2009). As the method used by Keubler et al (2009) was seen to be the most conservative (see Supplementary File) we focus on these results in the main text. Under this method, a glomerulus is considered to be a MG if its volume is greater than the 90th percentile of glomeruli volumes plus *k* times the difference between the 10^th^ and 90^th^ percentiles (Kuebler et al. 2010). We discriminated between MGs and normal glomeruli using a threshold k value of 1.5, which defines a moderate outlier (Sachs, 1988). MGs were considered to be present if they passed the discrimination threshold in both fully segmented males. We note that our definition of a MG is dependent upon the volumetric distribution of the other glomeruli, meaning that expansion of non-MGC glomeruli may obscure the presence of glomeruli expanded to a similar degree as MGs in other genera. We therefore provide the glomeruli size distributions for all species Figures S2-14.

For subsequent evaluations of sexual dimorphism we segmented: i) all glomeruli comprising the putative MGC, including both MGs and closely associated satellite glomeruli identified based on their physical position and internal structure, ii) the combined volume of all glomeruli, iii) total AL volume including glomeruli and the internal antennal lobe hub. Data for *Godyris zavalata* was taken from Montgomery and Ott (2015) but images were checked for consistency with newly obtained data.

### iv) Statistical analyses

In each species, linear models were used to test whether sexual dimorphism is present in the MGC glomeruli using the total volume of non-MGC glomeruli, calculated by subtracting the volume of MGC glomeruli from the total glomerular volume, as an additional factor to control for overall size. We also examined sexual dimorphism in total glomerular volume minus MGC volumes, correcting for the AL hub volume, a region of low synaptic density that is composed of tracts interlinking glomeruli and higher brain areas, to account for size variation. Normality of the residual errors was assessed using a Shapiro-Wilks test, and equal variance a Breusch-Pagan test. Data was seen to be normal in all cases. Where variance of errors was not equal we reassessed the data with robust error linear models. These corroborated the results of the linear models in all cases. Multiple testing was accounted for using a sequential Bonferroni correction (Benhamini and Hochberg, 1995). As sample size is limited in some species, we also report the Hedge’s *g* effect size (Hedges and Olkin, 1985) for all statistical tests.

## Results

### i) Ithomiines vary in the presence and form of the macro-glomerular complex

Ithomiine ALs are similar in structure to other Lepidoptera, glomeruli (averaging between 64-74 glomeruli; Table S2) positioned around a central fibrous region, the ‘AL hub’, which contains projections between glomeruli and downstream targets (Huetteroth and Schachtner, 2005; El Jundi et al., 2009b; Heinze and Reppert, 2012; Montgomery and Ott, 2015). MGs that passed our statistical threshold were observed in *Melinaea, Mechanitis, Forbestra, Ithomia, Hypothryris, Pseudoscada*, and *Hypoleria* (Figure 1). These structures were all components of a multi-glomerular complex (MGC) located at the dorsal base of the antennal nerve, with the exception of *P. florula* where we observe one MG within the MGC, and a second on the dorsomedial AL surface. Two MGs within the putative MGC cluster are observed in *Mechanitis* and *Melinaea* (Table S2, Figure S3-S4). In species lacking MGs, with the exception of *Methona*, we still identify a glomerular complex at the dorsal base of the antennal nerve that is distinct from other glomeruli in the remaining four species (Figure 1; Figure S2-4, 6-14). Despite the variable presence of a formally determined MG within this structure, we refer to it as an MGC throughout as we hypothesise it has a homologous function across species, with differential expansion of individual glomeruli.

**Figure One:**
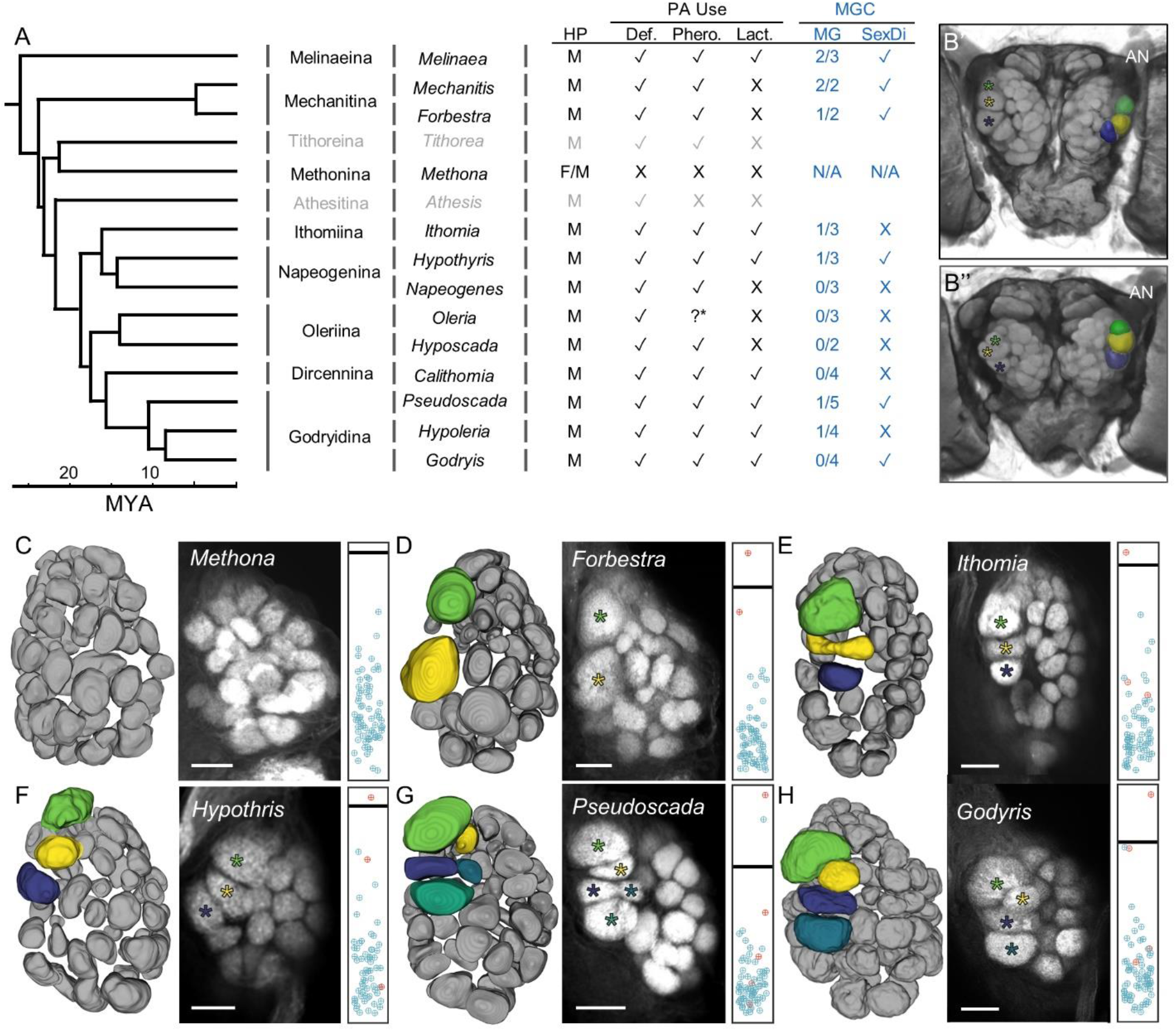
**A)** Ithomiine phylogeny illustrating variation in PA usage and summarizing results on MGC morphology. The phylogeny is based upon Chazot et al. 2019, and displays ithomiini subtribes and representative genera (Browers et al. 2014) with associated PA usage and putative MGC data. Subtribes where MGC was not evaluated are shown in grey. HP denotes presence of absence of hairpencils, and the sex where these are presence is indicated by F/M for female and male. PA use is categorised into three primary categories, use of PAs for defense, PAs as pheromones, and whether these PA pheromones contain lactones, Def,, Phero., and Lact., respectively. MGC information is given for number of MGs and total MGC glomeruli (MG) and whether there is evidence of sexual dimorphism in any MGC glomeruli (SD). **B)** Volume rendering of synapsin (3C11) immunofluorescence depicting brain anatomy, highlighting the position and morphology of MGC highlighted in colour, in *Oleria gunilla* **B’)** and **B”)** *Napeogenes larina*. **C-H)** Surface model of full glomeruli segmentations, an example of anti-synapsin immunofluorescence in an antennal lobe confocal sections, and the distribution of glomerular volumes in illustrative genera. In glomerular distribution plots, the discrimination threshold above which a glomerulus is considered a MG is indicated in black, and glomeruli within the putative MGCs highlighted in red. The genera shown are *Methona, Forbestra, Ithomia, Hypothyris, Pseudoscada*, and *Godyris*, **C-H)** respectively. For full results see Figure S2-14.

These MGCs occur in a corresponding position to that observed in *Godyris*, but the degree to which this structure is raised with respect to the AL surface is variable. Oleriina species exhibit a more homogenous glomerular surface than those of other subtribes, whereas the MGC is particularly pronounced in species of Dircennina and Godyridina (Figure S11-14). The composition of the MGC is also highly variable across the tribe. In the basal Mechanitina, the MGC contains two glomeruli. In representatives of Melinaeina, Ithomiina, Napeogenina, and Olerinia we see a single additional satellite glomerulus. Despite being within *Olerinia, Hyposcada* does not share this satellite glomeruli, and the two observed MGC glomeruli are of reduced size. The MGC observed in Dircennina and Godyridina have acquired further satellite glomeruli, with their MGCs containing 4 glomeruli, or 5 in the case of *Pseudoscada*. The most morphologically complex MGC is observed in *Pseudoscada*, with one MG and four satellite glomeruli.

### ii) Variation in sexual dimorphism in antennal lobe structures

Within the MGC, we also observe varying levels of sexual dimorphism across ithomiines. The size of at least one glomerulus is sexually dimorphic, being of greater size in males, in *Mechanitis, Forbestra*, *Hypothyris*, *Pseudoscada*, and *Godyris* (Table 1a). This is limited to a single glomerulus in all genera with the exception of *Pseudoscada*, which has three sexually dimorphic glomeruli (Table 1a; Figure 2). After multiple test correction, we do not observe dimorphism in any MGC glomeruli of *Melinaea, Ithomia, Napeogenes, Hyposcada, Oleria, Calithomia, and Hypoleria* (Table 1a). However, we note that in many non-significant tests we observe effect sizes that are comparable with those observed in sexually dimorphic genera (Table 1a) (Cohen, 1988). This may suggest that with increased sample size some of these genera would also display significant levels of sexual dimorphism. The total volume of non-MGC glomeruli were seen to be sexually monomorphic, with the exception of *Hyposcada* (Table 1b).

**Table 1a:**
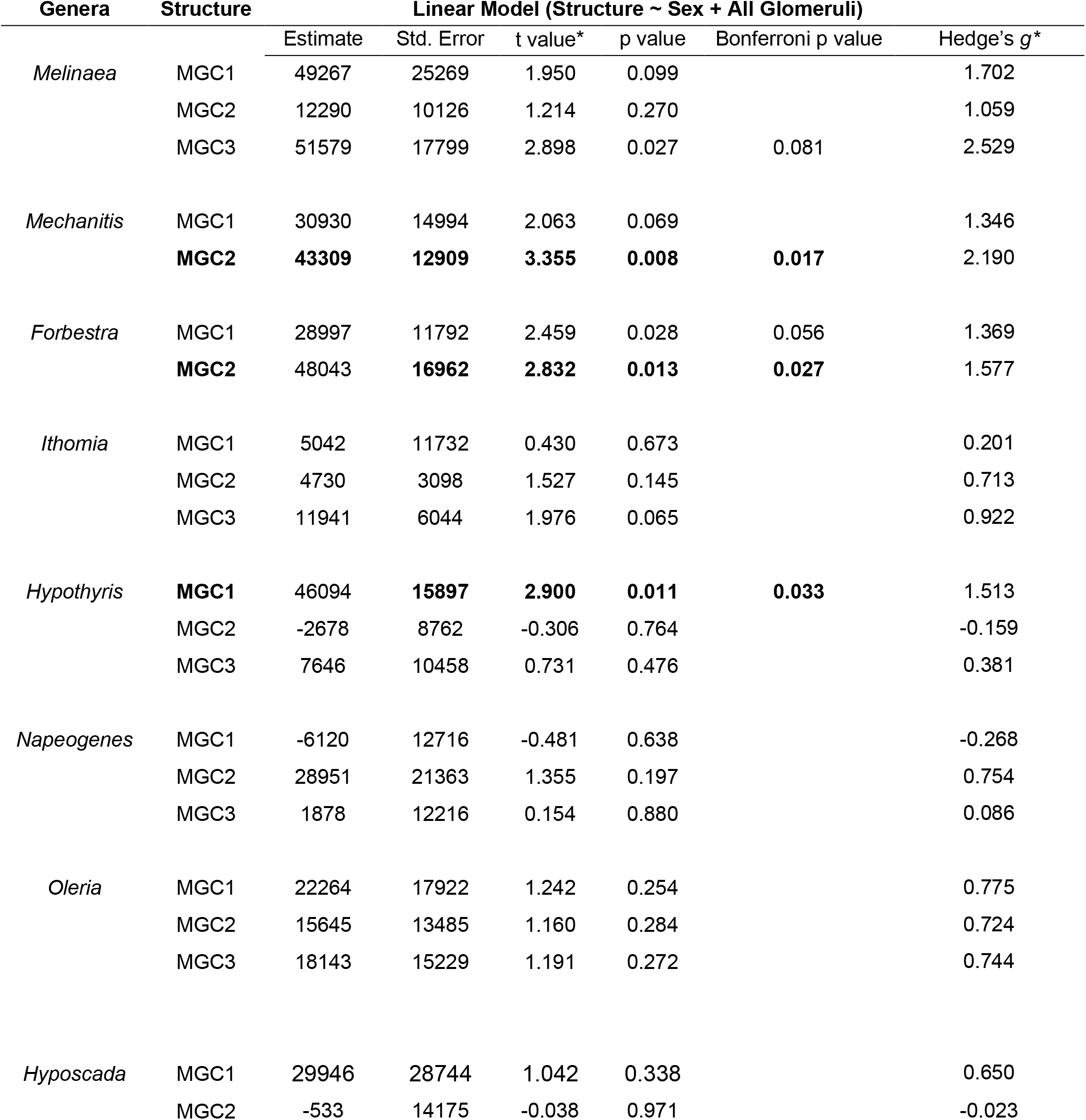
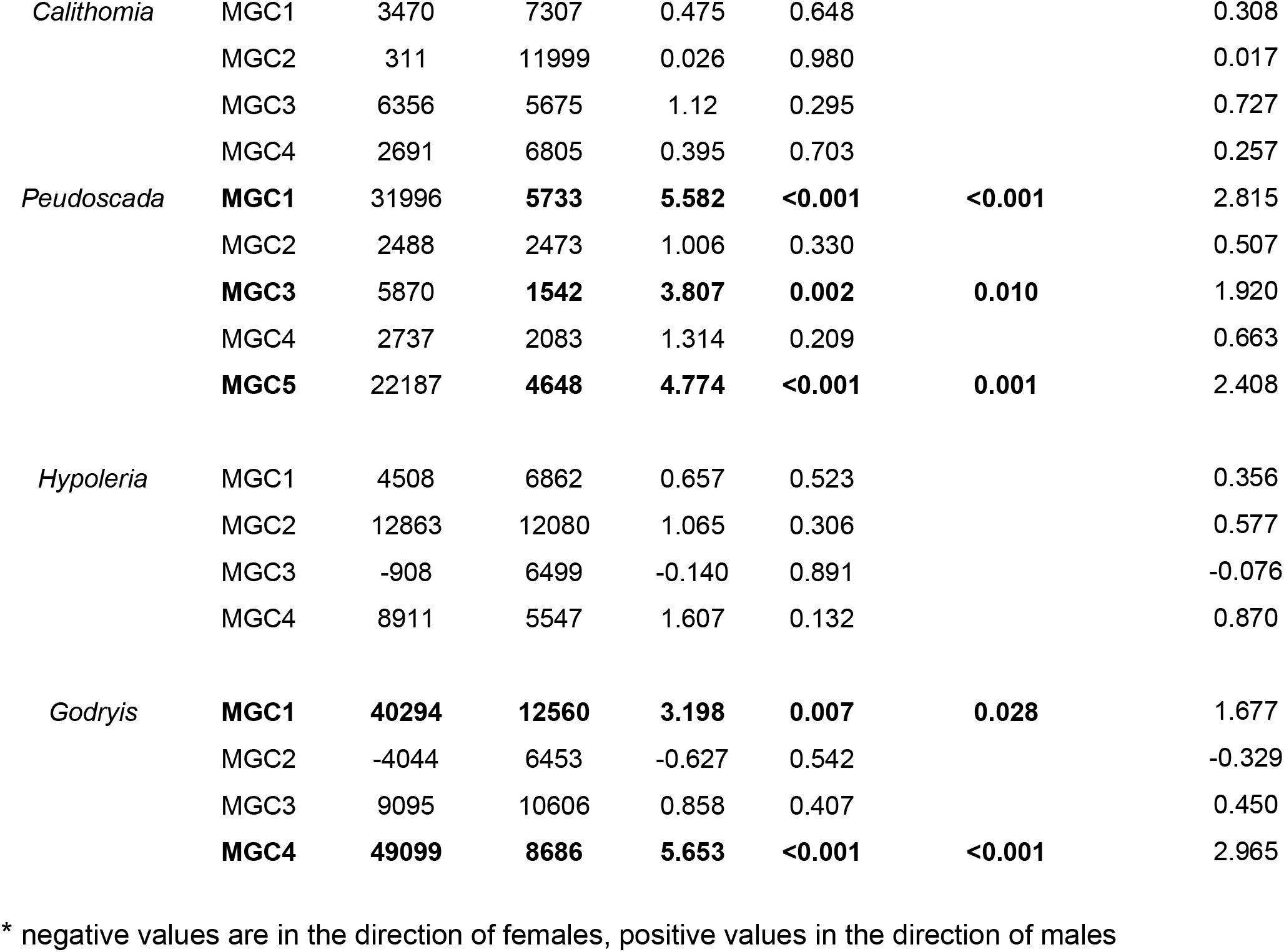
Statistical tests for MGC glomeruli sexual dimorphism, evaluating the influence of sex with the total non-MGC glomeruli volume as a second factor. Significant results are highlighted in bold.

**Table 2:** Results of Statistical tests AL sexual dimorphism, evaluating the influence of sex with the AL-hub volume as a second factor. Significant results highlighted in bold.

| Genera | Linear Model (AL ~ Sex + AL Hub) |  |  |  |  |
| --- | --- | --- | --- | --- | --- |
| | Estimate | Std. Error | t value* | p value | Hedge's $g^*$ |
| *Melinaea* | -22946 | 844708 | -0.027 | 0.979 | -0.024 |
| *Mechanitis* | 238652 | 487445 | 0.490 | 0.636 | 0.320 |
| *Forbestra* | 361659 | 278020 | 1.301 | 0.214 | 0.724 |
| *Ithomia* | 73192 | 246454 | 0.297 | 0.770 | 0.139 |
| *Hypothyris* | -104491 | 473039 | -0.221 | 0.828 | -0.115 |
| *Napeogenes* | 157416 | 316589 | 0.497 | 0.627 | 0.277 |
| *Oleria* | -63231 | 229058 | -0.276 | 0.790 | -0.172 |
| ***Hyposcada*** | **-1377952** | **315420** | **-4.369** | **0.005** | **-2.727** |
| *Calithomia* | -450012 | 275856 | -1.631 | 0.141 | -1.059 |
| *Peudoscada* | 67279 | 149040 | 0.451 | 0.658 | 0.228 |
| *Hypoleria* | -481089 | 376968 | -1.276 | 0.224 | -0.691 |
| *Godryis* | 57206 | 279056 | 0.205 | 0.841 | 0.108 |

\* negative values are in the direction of females, positive values in the direction of males

**Figure Two:**
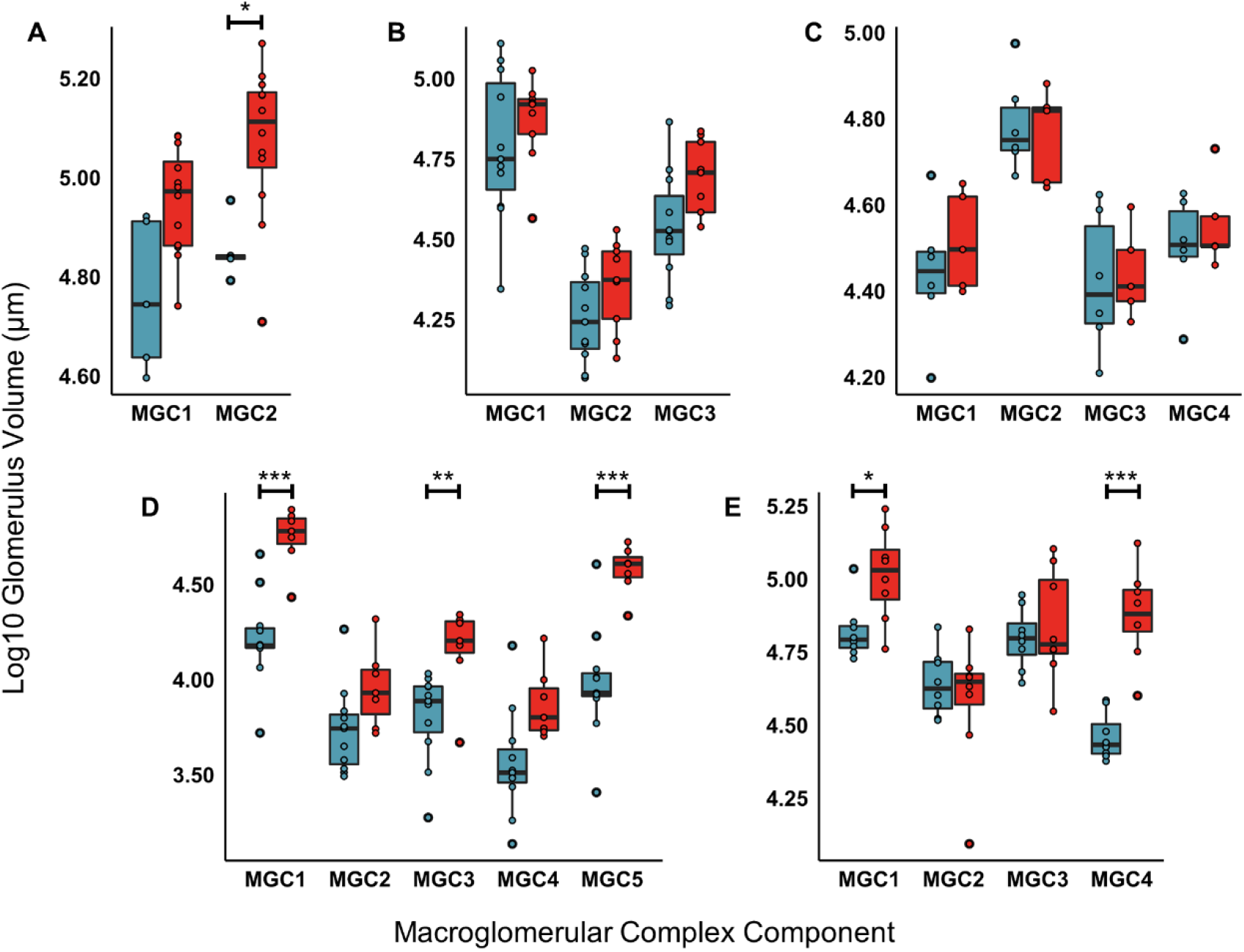
Varying levels of sexual dimorphism observed in MGC glomeruli volumes across selected ithomiine genera showing (A-E) *Forbestra* (♀5, ♂12), *Ithomia (*♀11, ♂9), *Callithomia (*♀5, ♂6), *Pseudoscada (*♀11, ♂7), *Godyris (*♀8, ♂8), respectively. Females are shown in blue, males are shown in red. Significance denoted by: *<0.05, **<0.01, ***<0.001. For full results see Figure S2-14.

## Discussion

Within Lepidoptera, significant variation has been observed in the volume of olfactory processing areas. In moths, males have vastly expanded glomeruli in the antennal lobe (MGs/MGCs) that are typically sexually dimorphic and responsive to pheromones (Bretschneider, 1924; Matsumoto and Hildebrand, 1981; Koontz and Schneider, 1987). Here, we present evidence for the widespread distribution of MGCs across the Ithomiini, the only tribe of butterflies currently known to possess this kind of olfactory specialisation (Montgomery and Ott, 2015). We provide evidence that, having secondarily re-evolved this AL specialisation, ithomiine MGC have diversified, and are variable across species in both composition and the degree of sexual dimorphism.

Two hypotheses have been proposed to explain why ithomiines have secondarily acquired a trait lost at the base of butterflies. First, ithomiine MGCs could reflect heightened sensitivity of males to PA resources, increasing their foraging efficiency (Montgomery and Ott, 2015). PAs are plant-derived chemicals utilised for several key fitness traits; chemical defence, pheromone synthesis, and nuptial gifts (Pliske, 1975, Pliske et al. 1976; Brown, 1984). Scarcity of PA resources, or competition amongst males, may have increased the selective pressure for sensitivity to PAs and led to expanded processing regions. This hypothesis would predict a consistent pattern of MG presence and sexual dimorphism across all species with this ecological strategy. However, our results are not consistent with this prediction. Both the presence of an MG and the presence and degree of sexual dimorphism vary across ithomiine genera strongly attracted to PAs. This implies a lack of *direct* association between PA use, or detection, and MGCs. Despite its variability, a MGC, or at least a positionally homologous glomerular cluster, is identifiable in all members of the tribe, with the exception of *Methona*. The absence of an MGC in *Methona* may instead be consistent with an *indirect* relationship between the use of PAs and the presence of an MGC, as this genus does not strongly rely on PAs for chemical defence, nuptial gifts or pheromone precursors (Brown, 1987; McClure et al. 2019). In addition, representatives of subtribes with the most morphologically pronounces MGCs (Godyriina, Ithomiia) have particularly strong male responses to PA baits (Pliske 1975, Brown 1984). Despite this, the lack of ubiquitous sexual dimorphism and inconsistency of MGC structure argue against a singular role for the MGC in PA foraging. Hence, the structural diversification of the ithomiine MGC may relate to *how*, rather than merely *if*, PAs are used in sexual communication.

Indeed, the second hypothesis for the origin of this structure suggests a role in pheromone processing (Montgomery and Ott, 2015). Across ithomiines, male pheromones are used for the attraction of females, with some lineages additionally utilising pheromones in male territorial defence (Pliske 1975). In many of these species, males are repulsed by lactones, which are present in the pheromone blend of phylogenetically disparate ithomiines (Schulz et al. 2004). Genera whose pheromone blends contain lactones (*Melinaea, Ithomia, Hypothyris, Calithomia, Pseudoscada, Godyris*, and *Hypoleria* (Schulz et al. 2004)) all have additional glomeruli incorporated into their MGCs. Whilst expanded MGCs are also observed in *Napeogenes* and *Oleria*, which lack pheromonal lactones, these genera also have novel pheromone blends, utilising compounds that are not observed in other ithomiines (Shultz et al. 2004; Stamm et al. 2019). The differentiation of pheromone repertoires in these species may suggest that OR duplication events have enabled the utilisation of additional pheromones, that were either latent or thereafter derived. In addition, in genera where the expression of PA-derived pheromones is reduced, such as *Hyposcada* (Schulz et al. 2004; Stamm et al. 2019), we observe a reduction in the volume and/or number of MGC glomeruli. These features suggest a significant role of pheromone usage in the function of the ithomiine MGC. The position of the ithomiine MGC is also comparable to the putative pheromone processing cluster of glomeruli in *D. plexxipus*, which shares a similar fibrous structure (Heinze and Reppert, 2011; Montgomery and Ott, 2015). The important role that PAs play in the ecology of Danaid species may suggest that this glomerulus was ancestrally sensitive to PA-derived chemicals in Danaids and ithomiines, and was elaborated in ithomiines (Brown 1987, Schulz et al. 1988).

While our data is consistent with a role for the MGC in sexual communication, sexual dimorphism is variable across the tribe, showing no clear phylogenetic pattern. The lack of consistent sexual dimorphism may reflect similar reliance of each sex on these cues, but in differing contexts and/or differing valances. PA-derived pheromones have been hypothesised to represent honest signals of male quality (Trigo et al. 1994), which may be salient cues for both reproductively receptive females, and males during male-male competition. In this case sexual dimorphism could be explained by inhibitory adaptation, in which chronic exposure to odorants results in the expansion of responsive glomeruli through increased innervation by inhibitory AL local neurons (Devaud et al. 2001; Sachse et al. 2007; Anton et al. 2015). Male ithomiines are strongly attracted to PA resources, with sensitivity to these compounds essential, despite being a source of these volatile compounds themselves (Brown, 1984; Trigo et al. 1996). Male-biased sexual dimorphism may therefore be a mechanism to overcome chronic self-exposure to PAs. This hypothesis would predict varying degrees of sexual dimorphism depending on both the amount of pheromone utilised by males, and the similarity of compounds in the male-pheromone blend and PAs used for chemical defence. Further, we note that as our samples are derived from wild populations and some of our volumetric estimates show high levels of intraspecific variation, which could be consistent with a plastic response to differential exposure to PAs/PA-derivatives in the adult or juvenile environment.

While neither hypothesis solely explains the emergence and full diversity of ithomiine MGCs, the combination of selection associated with finding and utilising PAs for pheromonal compounds likely explains both for the origin and diversification of this structure, respectively. However, the mechanism behind this neural innovation is unclear. Butterfly pheromones are processed by glomeruli that are also responsive to plant odours(Larsdotter-Mellström et al. 2016), unlike moths, in which pheromone signals are processed by a separate subset of glomeruli from other odours (Kanzaki and Shibuya, 1983; Christensen and Hildebrand, 1987; Hansson et al. 1991). The evolution of a MGC in the evolutionary context in which the pheromone response pathway is integrated into general odour processing raises several questions: i) did the MGC originate from existing glomeruli that were sensitive to plant emitted PAs, becoming specialised to detect PA-derived pheromones? ii) were MGC glomeruli, and their associated olfactory receptors, produced by co-option of existing receptors or duplication of PA-sensitive receptors? iii) do pheromonal responses remain integrated with general odours during olfactory processing or have parallel pathways evolved in ithomiines?

We suggest that these questions can be answered through a combination of functional assays and phylogenetics. For example, the response of MGs to pheromone extracts and/or plant PAs can be assessed using electrophysiology or in vivo Ca^2+^ imaging, in ithomiines and in danaids comparing putatively homologous glomeruli. Similar approaches can be used to evoke odour maps in response to different stimuli (Galizia et al. 2000; Carlsson et al. 2011) which would demonstrate whether pheromones are encoded in combination with general odours, as observed in other butterflies (Larsdotter-Mellström et al. 2016), or separately, as seen in moths (Boeckh et al. 1965). Finally, identifying ORs that are functionally linked to MGCs through analyses of their sex-biased expression in species showing glomerular dimorphism, combined with phylogenetic studies to test homology of these loci with plant-detecting or pheromone-detecting ORs in other Danaiids and more broadly across Lepidopterans, would provide a test of the ancestral function of this pathway.

In conclusion, we have identified a highly dynamic cluster of glomeruli in the ithomiine tribe that includes significantly expanded MGs. To date, ithomiines are the only tribe of butterflies reported to have acquired this AL specialisation, strongly implying it has evolved secondarily. We show that ithomiine MGCs are variable in presence, composition, and degree of sexual dimorphism. Our data supports the hypothesis that foraging for plant derived chemical defence may be the ancestral source of selection pressure favouring the evolution of MGCs in this tribe, with their subsequent elaboration associated with the diversification of ithomiine pheromonal cues. This line of communication could be particularly important in ithomiines as they form multi-species mimicry rings, with ecological convergence within co-mimetic species (Elias et al. 2008) potentially rendering long range visual mating cues less reliable.

## Acknowledgements

We thank Álvaro Barragán, Emilia Moreno, PabloJarrín, and David Lasso from the Estación Científica Yasuní and Pontificia Universidad Católica del Ecuador, and Maria Arévalo from the Parque Nacional Yasuní Ministerio Del Ambiente for assistance with collection and exportation permits, and Francisco Ramlrez Castro for assistance in the field in 2011. We also thank Swidbert Ott for advice early on in this project, and Matt Wayland and the Imaging Facility at the Dept of Zoology, University of Cambridge for confocal support. This work was funded by a Royal Commission for the Great Exhibition Research Fellowship, a Royal Society Research Grant and a NERC IRF (NE/N014936/1) to SHM.

## Notes

### Competing Interest Statement

The authors have declared no competing interest.

