## Supplementary material for "Re-emergence and diversification of a specialised antennal lobe morphology in ithomiine butterflies": Re-emergence and diversification of a specialised antennal lobe morphology Supplementary File.pdf

### Contents

1. Sampling, page 2
2. Macroglomerulus identification, page 3
3. Anatomical descriptions of each genus, page 5

### 1. Sampling

Species were chosen for phylogenetic coverage among samples with sufficient sample size. Where limited single species samples were available, in subtribes deemed important, samples were pooled within genera in order to reach a minimum sample size.

**Table S1:** Samples utilised in this study. Data from samples marked with * were taken from Montgomery and Ott, 2015.

| Subtribe | Genera | Species | Number of Samples | | |
|---|---|---|---|---|---|
| | | | MG Discrimination | Sexual Dimorphism | |
| | | | Males | Males | Females |
| Melinaeina | *Melinaea.* | *spp.* | 2 | 2 | 7 |
| | | *menophilus* | 0 | 0 | 3 |
| | | *mnasius* | 1 | 1 | 3 |
| | | *satevis* | 1 | 1 | 1 |
| Methonina | *Methona* | *spp.* | 4 | N/A | N/A |
| | | *grandior* | 2 | N/A | N/A |
| | | *curvifascia* | 2 | N/A | N/A |
| Mechanitina | *Mechanitis* | *polymnia* | 2 | 8 | 4 |
| | *Forbestra* | *olivencia* | 2 | 12 | 5 |
| Ithomiina | *Ithomia* | *amarilla* | 2 | 9 | 11 |
| Napeogenina | *Hypothyris* | *anastasia* | 2 | 12 | 6 |
| | *Napeogenes* | *larina* | 2 | 12 | 5 |
| Oleriina | *Oleria* | *gunilla* | 2 | 7 | 5 |
| | *Hyposcada* | *illinissa* | 2 | 7 | 5 |
| Dircennina | *Calithomia* | *lenea* | 2 | 6 | 5 |
| Godyridina | *Pseudoscada* | *florula* | 2 | 7 | 11 |
| | *Hypoleria* | *sarepta* | 2 | 10 | 6 |
| | *Godyris* * | *zavaleta** | 2* | 8* | 8* |

### 2. Macroglomerulus identification

We used three different techniques for the discrimination of macroglomeruli that have previously been applied in previous publications.

1) Montgomery and Ott (2015) - 2 Standard Deviations

Glomeruli are considered macroglomeruli if their absolute volume exceeds that of the mean plus two times the standard deviation, i.e.:

$$x > \bar{x} + (2 \times S.D)$$

Where: x = focal glomerular volume, $\bar{x}$ = glomerular volume mean, S.D = standard deviation.

2) Keubler et al. (2010) - K-Variance

A macroglomeruli has a volume that is greater than the volume of the 90th percentile of glomeruli volumes plus k times 80% of the observed volume, i.e.:

$$x > x^{90} + k(x^{90} - x^{10})$$

Where: x = glomeruli volume, $x^{90}$ = the volume of the 90% percentile glomeruli, $x^{10}$ = the volume of the 10% percentile glomeruli, and k = threshold variable.

The threshold value used in the present paper is 1.5. Whilst this is lower than the k value of 3 that Keubler et al. used, the authors noted that their value was highly conservative (Keubler et al. 2010), an effect which is likely exacerbated in species with smaller numbers of glomeruli in lepidoptera. In addition, the distribution of the data in the original paper (Keubler et al. 2010) is narrower than the data presented here. Our *k* value defines a moderate outlier (Sachs, 1988).

3) Kelber et al. 2009 - Radial

This method transforms the volumetric values in order to account for non-normality in the distribution of glomerular volumes. It assumes that glomerular volumes are perfect spheres, and the radius value of each is taken as a measure of size. The mean radius and the standard deviation of the radii are then calculated. Each glomerulus is assigned the value of its radius minus that of the mean radius, divided by the standard deviation of radii. If this final value is greater than a defined cut-off value it is deemed a macroglomerulus, i.e.:

$$\frac{(x_r - \bar{x}_r)}{\text{S.D.}} = x_v \ , \ x_v > \text{CV}$$

Where: $x_r$ = radius of sphere with volume x, $\bar{x}_r$ = mean radius calculated from all glomeruli, S.D. = standard deviation of calculated radii, and CV = cut-off value.

This can be rearranged to give a cut-off value in $\mu m^2$. Whilst this affects the distribution of the data with respect to the cut off value, it does not affect the discrimination between macroglomeruli and glomeruli itself.

$$\text{CV } \mu m^2 = \pi((CV \times SD) + \bar{x}_r)^2$$

For our cut-off value we used a value of 3. This cut-off value is lower than the highly conservative value used originally but is justified due to the lower number of glomeruli that are observed in Lepidoptera vs Hymenoptera, and a narrower distribution of the data observed than in the original paper (Kelber et al. 2010).

Table S2 provides the results of all three methods. As the method of Keubler et al. 2010 proves the most conservative in the majority of samples, we have used this method for the basis of our results in the main text. In addition, putative macroglomeruli were required to pass this threshold in all of the evaluated samples.

**Table S2:** Macroglomerulus identification using alternative methods

| Genera | Species | Total Glomeruli | Number of Macroglomeruli | | |
|---|---|---|---|---|---|
| | | | 2.S.D | K-Var | Radial |
| *Melinaea* | *mnasius* | 72 | 3 | 3 | 3 |
| | *satevis* | 76 | 2 | 2 | 2 |
| *Methona* | *curvifascia* | 70 | 2 | 0 | 0 |
| | | 74 | 3 | 0 | 1 |
| | *grandior* | 71 | 2 | 0 | 0 |
| | | 68 | 2 | 0 | 0 |
| *Mechanitis* | *polymnia* | 67 | 2 | 2 | 2 |
| | | 64 | 2 | 2 | 2 |
| *Forbestra* | *olivencia* | 66 | 2 | 2 | 2 |
| | | 65 | 2 | 1 | 1 |
| *Ithomia* | *amarilla* | 68 | 4 | 1 | 1 |
| | | 71 | 2 | 1 | 1 |
| *Napeogenes* | *larina* | 69 | 2 | 1 | 1 |
| | | 75 | 4 | 0 | 0 |
| *Hypothyris* | *anastasia* | 67 | 2 | 2 | 2 |
| | | 63 | 3 | 1 | 1 |
| *Hyposcada* | *illinissa* | 69 | 3 | 1 | 1 |
| | | 68 | 4 | 0 | 0 |
| *Oleria* | *gunilla* | 61 | 1 | 0 | 1 |
| | | 66 | 2 | 1 | 1 |
| *Calithomia* | *lenea* | 71 | 4 | 1 | 1 |
| | | 70 | 3 | 0 | 0 |
| *Pseudoscada* | *florula* | 73 | 2 | 2 | 2 |
| | | 72 | 5 | 4 | 3 |
| *Hypoleria* | *sarepta* | 73 | 3 | 1 | 2 |
| | | 74 | 3 | 1 | 1 |
| *Godrys* | *zavaleta* | 70 | 3 | 0 | 0 |
| | | 63 | 3 | 1 | 1 |

### 3. Anatomical descriptions of each genus

Species are listed in order of phylogeny displayed in Figure 1.

#### *i) Melinaea*:

We sampled 9 individuals from species *M.* menophilus, *M. mnasius*, and *M.* satevis (see Table S1). To investigate the interspecific variance in MGC component volumes across *Melinaea* species we performed a clustering analysis using hierarchical clustering with wards method. Volumes MGC component volumes were divided by the individual total glomerular volume, and then normalized by its row total (Figure S1). This shows that individual species do not cluster, with inter-individual variation being greater across our samples than interspecific. Given the limited samples size, we therefore grouped *Melinaea* species for subsequent analyses.

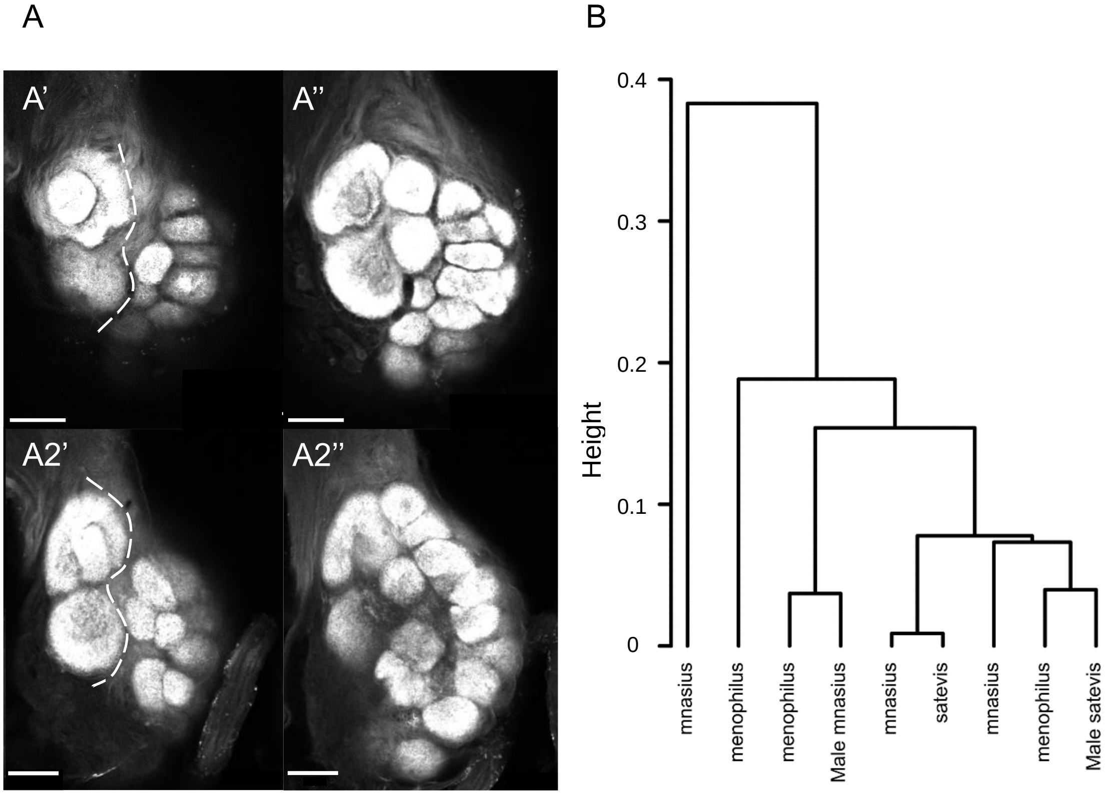


**Figure S1: A)** Confocal image of *Melinaea* ALs. MGC deliminated from rest of AL by dashed line. Scale bar = 50µm. **A1)** *Melinaea satevis* **A2)** *Melinaea mnanius*. Pannels depict surface of the antennal lobe and halfway through the MGC, **'** and **''** respectively. **B)** Hierarchical clustering of each *Melinaea* individual MGC, individuals are female unless otherwise indicated.

In the fully segmented males, 1 *M.* satevis and 1 *M.* mnasius, we observe two macroglomeruli (MGC1 and MGC3)(Figure S1 A, Figure S2 A), which are both part of a MGC (Figure S2 B, C) positioned at the base of the antennal nerve. MGCs across species showed no variation in the number of MGC components. We observe no statistically significant sexual dimorphism when controlling for variance in total glomerular volume (Figure S2 D, E).

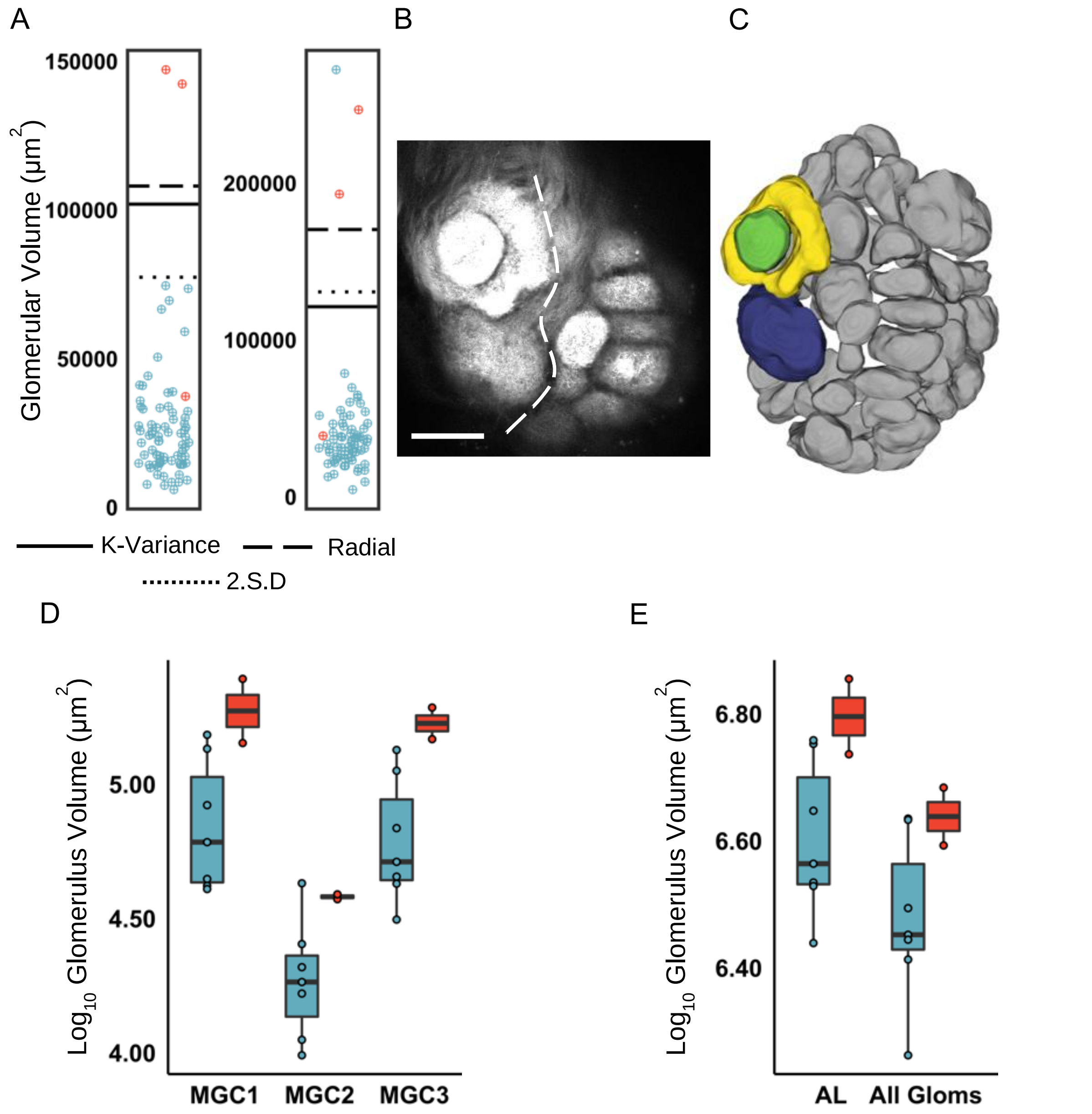


**Figure S2: *Melinaea MGC (♀7, ♂2)* A)** Scatterplot displaying individual glomerular volumes of two focal males. Glomeruli that are part of the putative MGC in Red, all other glomeruli Blue. Cut off lines indicated. **B)** Confocal image of *Melinaea* AL. MGC are deliminated from the rest of the AL by a dashed line. Scale bar = 50µm. **C)** Surface model showing segmented glomeruli from a focal male. MGC glomeruli coloured, other glomeruli in grey. **D)** Boxplot displaying volumes of female (blue) and male (red) MGC components. **E)** Boxplot showing volumes of AL and total non-MGC glomerular volumes of female (blue) and male (red). Significance from linear models shown by: *<0.05, **<0.01, ***<0.001.

#### *ii) Mechanitis*:

We sampled 4 male and 8 female *Mechanitis polymnia* in this study (see Table S1). We observe two macroglomeruli in *Mechanitis* (Figure S3 A), which constitute the MGC in this species (Figure S3 B, C). After correcting for multiple testing, we do not observe sexual dimorphism in either MG. However, we note the large effect sizes (Hedge's *d*: MGC1 = 1.346; MGC2 = 2.190).

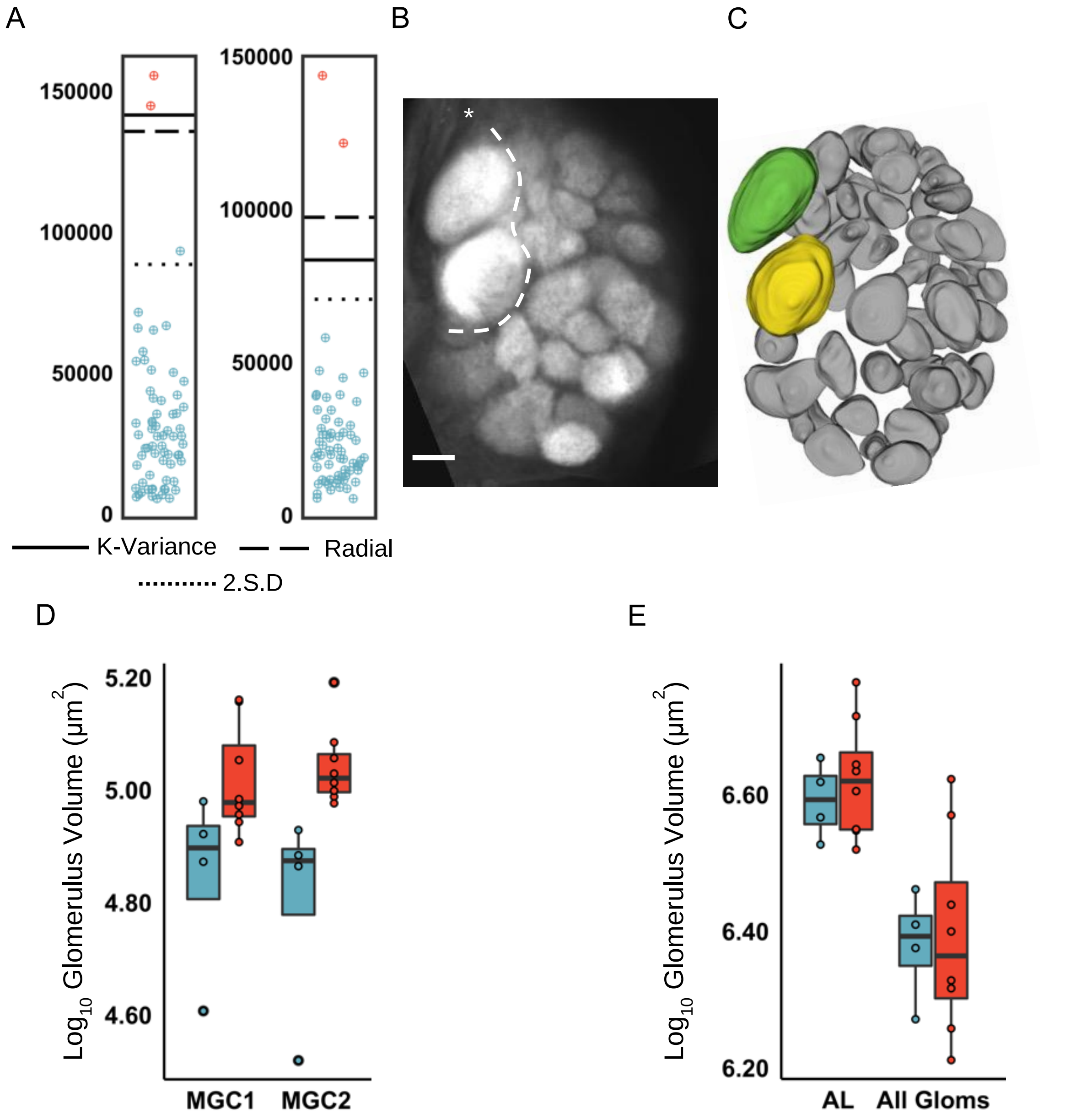


**Figure S3: *Mechanitis* MGC (♀4 ♂8) A)** Scatterplot displaying individual glomerular volumes of two focal males. Glomeruli that are part of the putative MGC in Red, all other glomeruli Blue. Cut off lines indicated. **B)** Confocal image of Mechanitis AL. MGC are deliminated from the rest of the AL by a dashed line. Scale bar = 50µm. **C)** Surface model showing segmented glomeruli from a focal male. MGC glomeruli coloured, other glomeruli in grey. **D)** Boxplot displaying volumes of female (blue) and male (red) MGC components. **E)** Boxplot showing volumes of AL and total non-MGC glomerular volumes of female (blue) and male (red). Significance from linear models shown by: *<0.05, **<0.01, ***<0.001.

### *iii) Forbestra*:

Twelve male and five female *Forbestra olivencia* were used to assess MGC presence, composition and sexual dimorphism. One MG is identified in *Forbestra* (MGC2) (Figure S4 A), as part of a MGC with an additional satellite glomeruli (Figure S4 B, C). After correcting for multiple testing, the MG (MGC2) is considered sexually dimorphic, however the satellite glomeruli does not pass our adjusted significance threshold. The degree of sexual dimorphism in MGC2 is pronounced (Hedge's *d*: MGC2 = 1.577). The effect size of sexual dimorphism in MGC1 is also large (Hedge's *d*: MGC1 = 1.369).

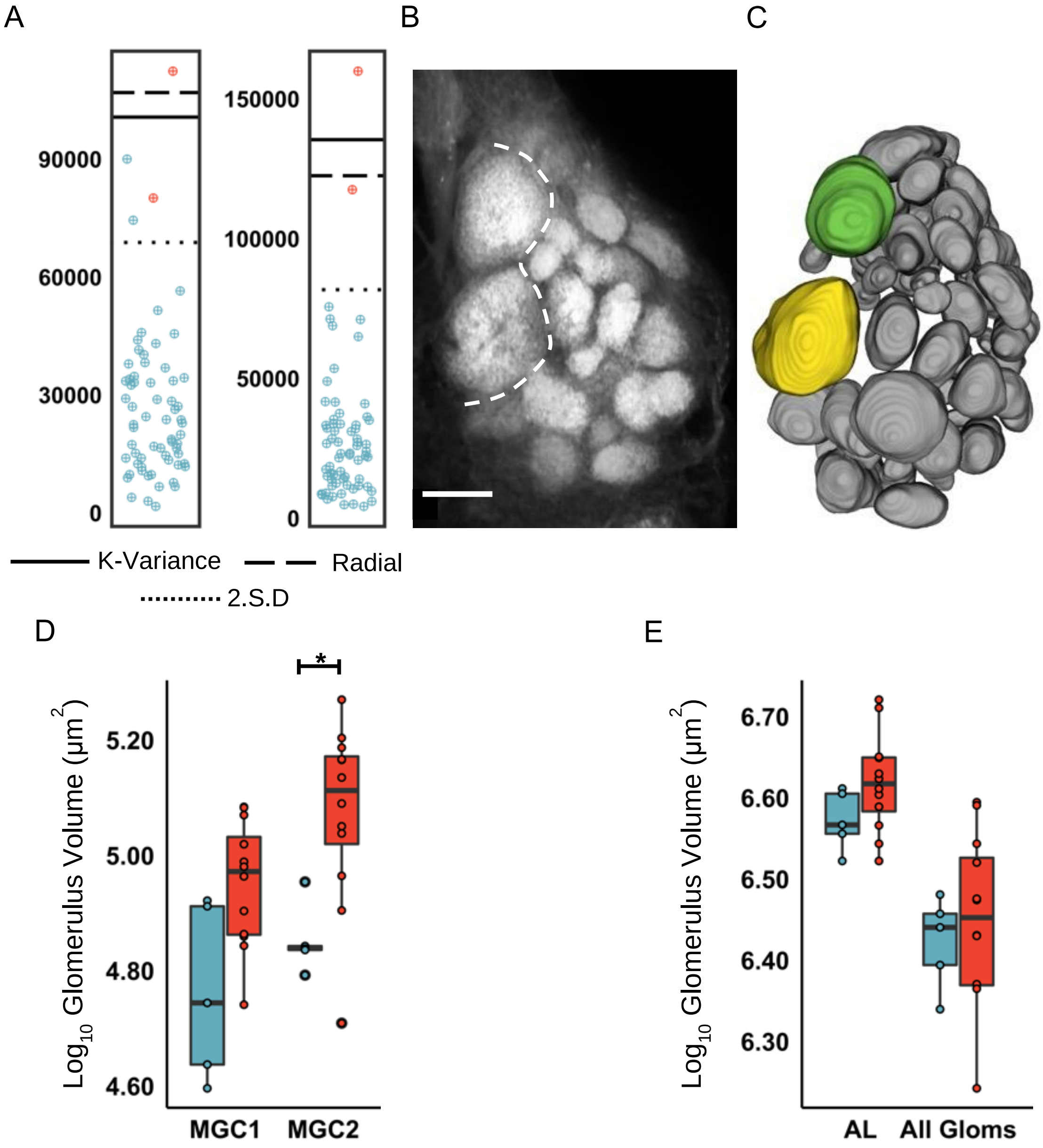


**Figure S4: *Forbestra* MGC (♀5, ♂12) A)** Scatterplot displaying individual glomerular volumes of two focal males. Glomeruli that are part of the putative MGC in Red, all other glomeruli Blue. Cut off lines indicated. **B)** Confocal image of *Forbestra* AL. MGC are deliminated from the rest of the AL by a dashed line. Scale bar = 50µm. **C)** Surface model showing segmented glomeruli from a focal male. MGC glomeruli coloured, other glomeruli in grey. **D)** Boxplot displaying volumes of female (blue) and male (red) MGC components. **E)** Boxplot showing volumes of AL and total non-MGC glomerular volumes of female (blue) and male (red). Significance from linear models shown by: *<0.05, **<0.01, ***<0.001.

### *iv) Methona*

We used two *Methona grandior* and two *Methona curvidascia* males to assess for the presence of an MGC in this genera. No glomeruli in either species reach our threshold for consideration as a MG in *Methona* (Figure S5 A, D). Additionally, we do not observe a glomerular cluster that is spatially homologous to the MGC seen in other ithomiine genera (Figure S5 B, C, E, F). Visual checks of female samples did not show obvious signs of a female specific MGC, though these were not quantitatively assessed.

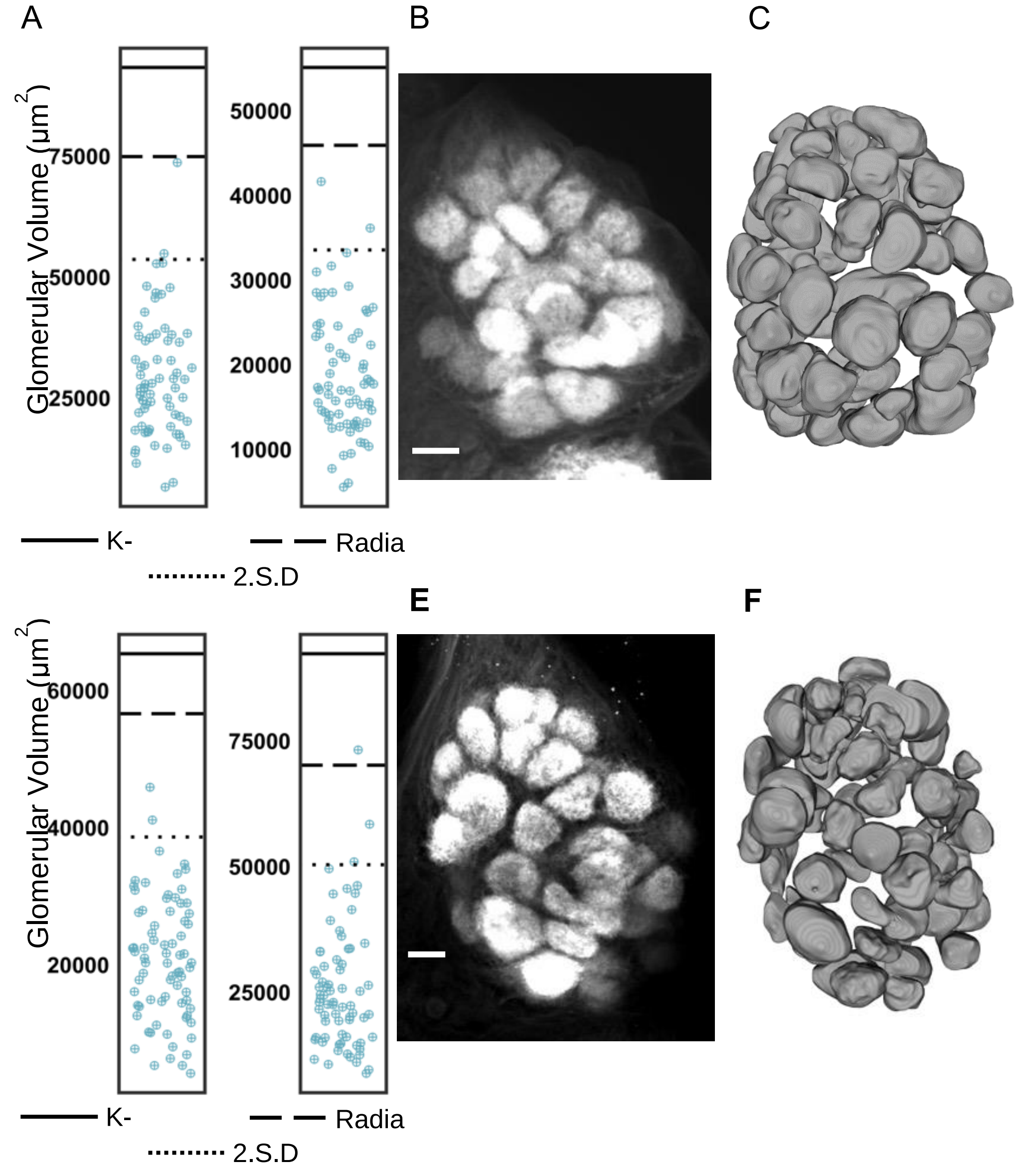


**Figure S5: Methona MGC (♂4) A)** Scatterplot displaying individual glomerular volumes of two *Methona grandior* males. Glomeruli that are part of the putative MGC in Red, all other glomeruli Blue. Cut off lines indicated. **B)** Confocal image of *Methona grandior* AL. Scale bar = 50μm. **C)** Surface model showing segmented glomeruli from a focal male. MGC glomeruli coloured, other glomeruli in grey. **D)** Scatterplot displaying individual glomerular volumes of two *Methona curvifascia* males. Glomeruli that are part of the putative MGC in Red, all other glomeruli Blue. Cut off lines indicated. **E)** Confocal image of *Methona curvifascia* AL. MGC deliminated from rest of AL by dashed line. Scale bar = 50μm. F**)** Surface model showing segmented glomeruli from a focal male. MGC glomeruli coloured, other glomeruli in grey. Significance from linear models shown by: *<0.05, **<0.01, ***<0.001.

### *v) Ithomia*:

Twenty *Ithomia amarilla* samples were used in this study, eleven females and nine males. A MGC composed of a single MG (MGC1) (Figure S6 A) and two satellite glomeruli (Figure S6 B, C) is identified. We note that MGC2 was separated into two glomeruli in a subset of individuals. Due to inconsistency in separation we have considered this as a single glomerulus in our analyses, however, it may be a case of two olfactory receptors that are in the process of diverging. We do not observe any sexual dimorphism is any glomeruli (Figure S6 D, E).

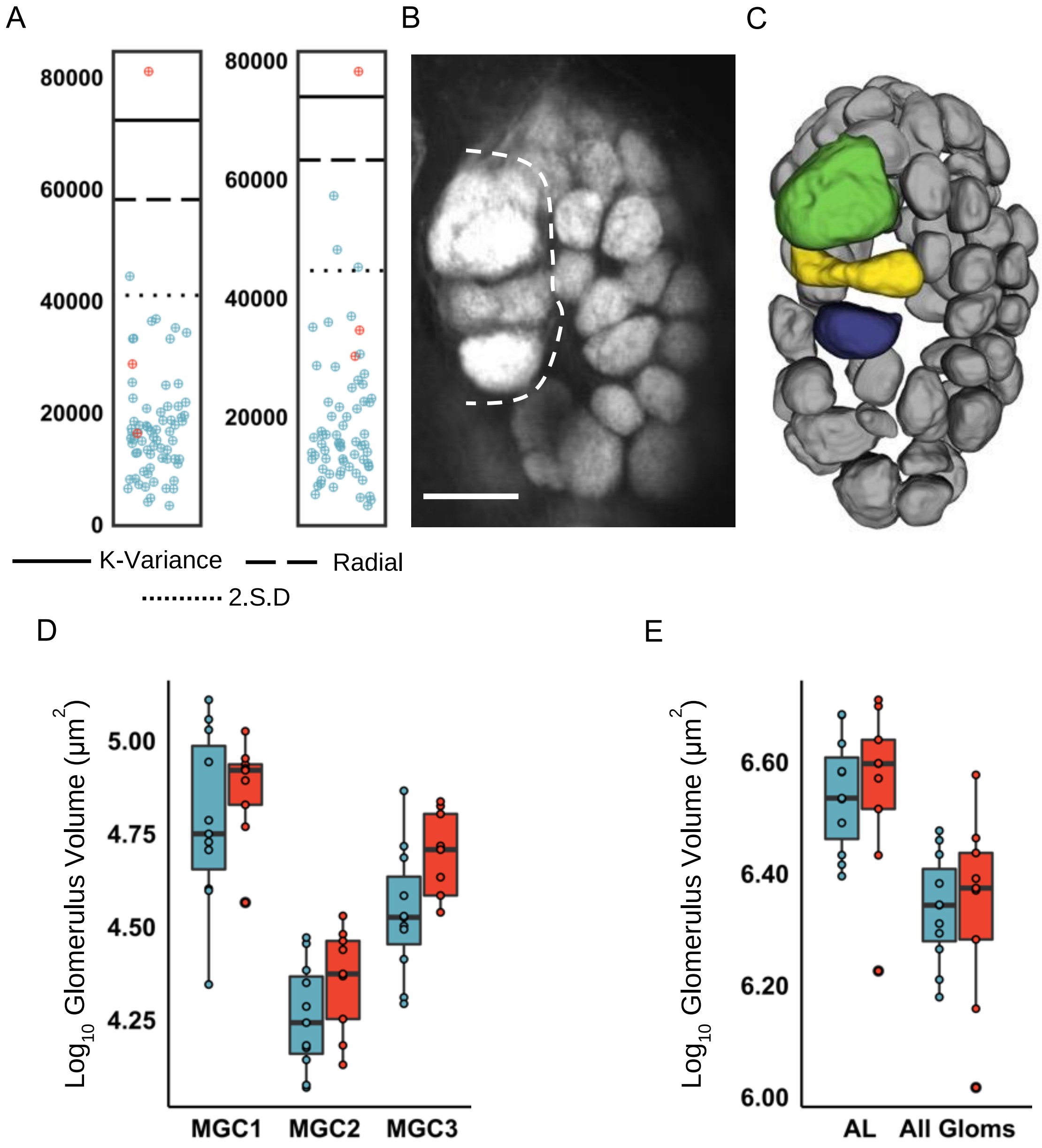


**Figure S6: *Ithomia* MGC (♀9, ♂11) A)** Scatterplot displaying individual glomerular volumes of two focal males. Glomeruli that are part of the putative MGC in Red, all other glomeruli Blue. Cut off lines indicated. **B)** Confocal image of *Ithomia* AL. MGC are deliminated from the rest of the AL by a dashed line. Scale bar = 50µm. **C)** Surface model showing segmented glomeruli from a focal male. MGC glomeruli coloured, other glomeruli in grey. **D)** Boxplot displaying volumes of female (blue) and male (red) MGC components. **E)** Boxplot showing volumes of AL and total non-MGC glomerular volumes of female (blue) and male (red). Significance from linear models shown by: *<0.05, **<0.01, ***<0.001.

### *vi) Hypothyris*:

Six female and twelve male *Hypothyris anastasia* were used in this study. We observe a single MG (MGC1) that is consistent across both evaluated individuals (Figure S7 A). Together with two satellite glomeruli, this forms a MGC (Figure S7 B, C). The MG (MGC1) is seen to be sexually dimorphic.

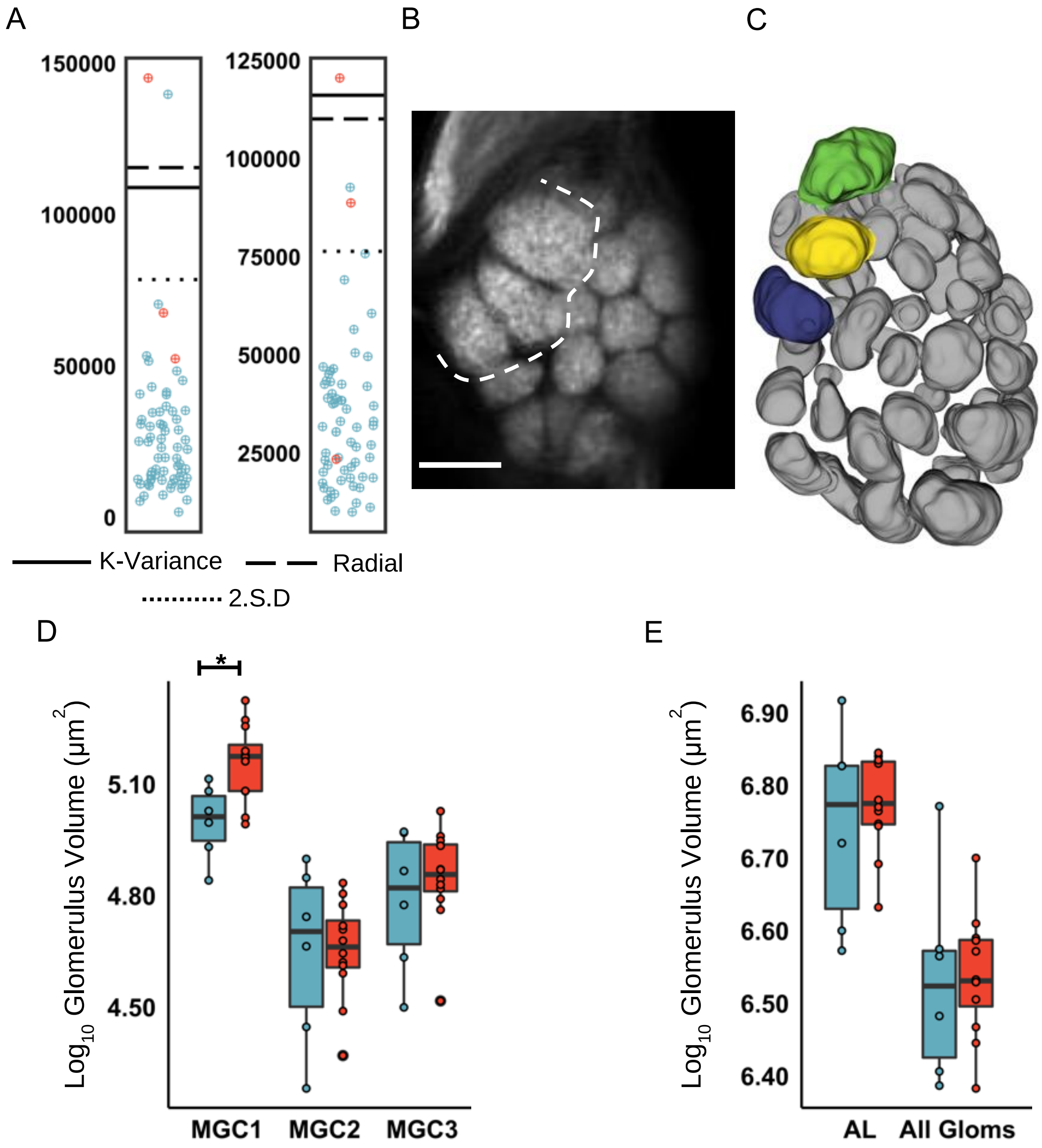


**Figure S7: *Hypothyris* MGC (♀6, ♂12) A)** Scatterplot displaying individual glomerular volumes of two focal males. Glomeruli that are part of the putative MGC in Red, all other glomeruli Blue. Cut off lines indicated. **B)** Confocal image of *Hypothyris* AL. MGC are deliminated from the rest of the AL by a dashed line. Scale bar = 50µm. **C)** Surface model showing segmented glomeruli from a focal male. MGC glomeruli coloured, other glomeruli in grey. **D)** Boxplot displaying volumes of female (blue) and male (red) MGC components. **E)** Boxplot showing volumes of AL and total non-MGC glomerular volumes of female (blue) and male (red). Significance from linear models shown by: *<0.05, **<0.01, ***<0.001.

**vii) *Napeogenes*:**

Seventeen *Napeogenes larina* individuals were sampled, twelve males and five females. We do not identify any MGs (Figure S8 A). However, we observe a glomerular cluster in a conserved position as the MGC of other ithomiines (Figure S8 B, C). None of the glomeruli of this spatially homologous cluster are sexually dimorphic. Glomerular volumes are seen to be especially variable in this genera.

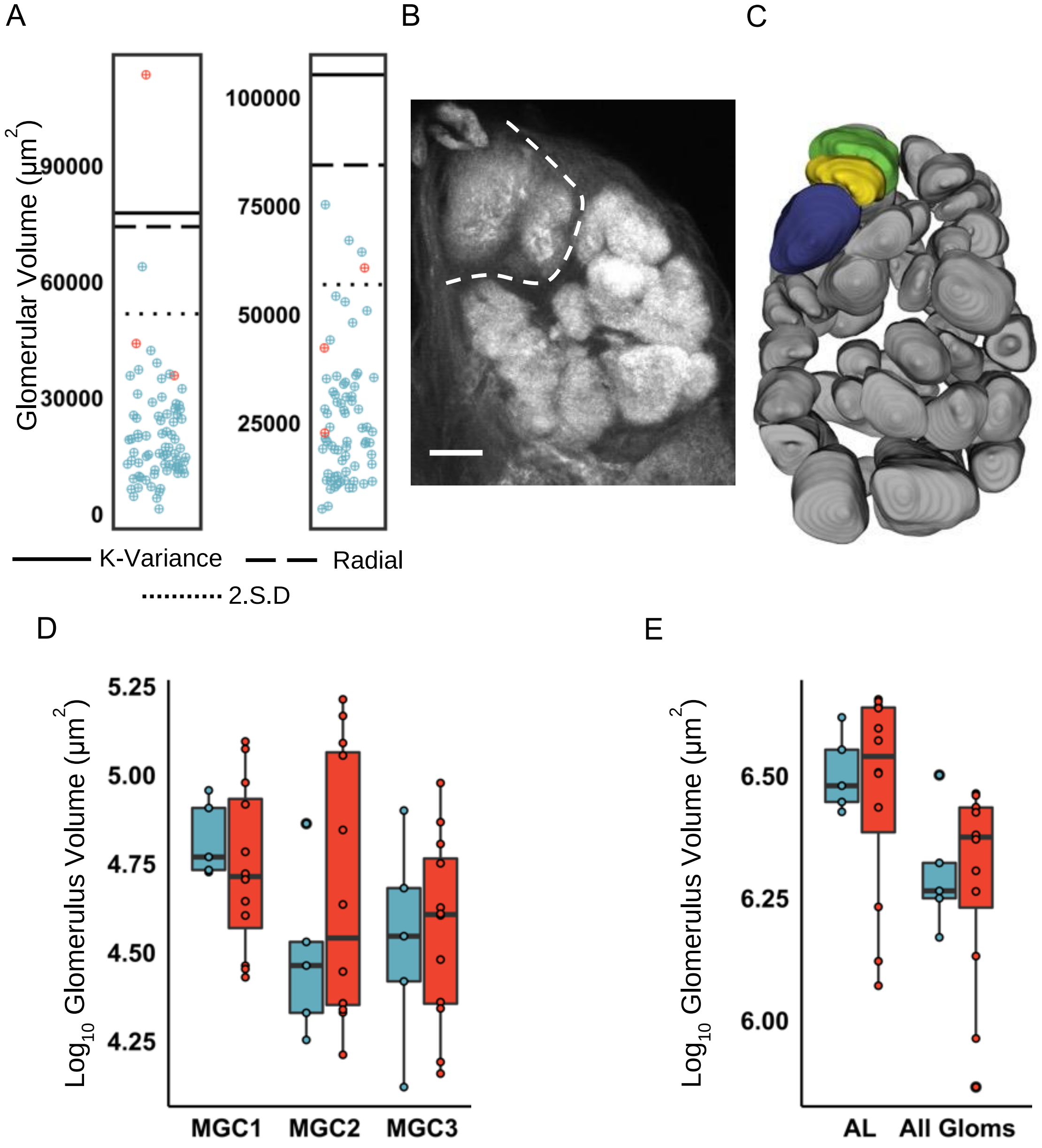


**Figure S8: *Napeogenes* MGC (♀5, ♂12) A)** Scatterplot displaying individual glomerular volumes of two focal males. Glomeruli that are part of the putative MGC in Red, all other glomeruli Blue. Cut off lines indicated. **B)** Confocal image of *Napeogenes* AL. MGC are deliminated from the rest of the AL by dashed line. Scale bar = 50µm. **C)** Surface model showing segmented glomeruli from a focal male. MGC glomeruli coloured, other glomeruli in grey. **D)** Boxplot displaying volumes of female (blue) and male (red) MGC components. **E)** Boxplot showing volumes of AL and total non-MGC glomerular volumes of female (blue) and male (red). Significance from linear models shown by: *<0.05, **<0.01, ***<0.001.

### ***viii) Oleria***:

Seven male and five female *Oleria gunilla* were used in this study. Whilst we do not observe a MG in this genera (Figure S9 A), a clearly distinct glomerular cluster, homologous in appearance to that of the MGC of other ithomiines, is identified (Figure S9 B, C). None of the components of this glomerular cluster are seen to be sexually dimorphic (Figure S9 D, E).

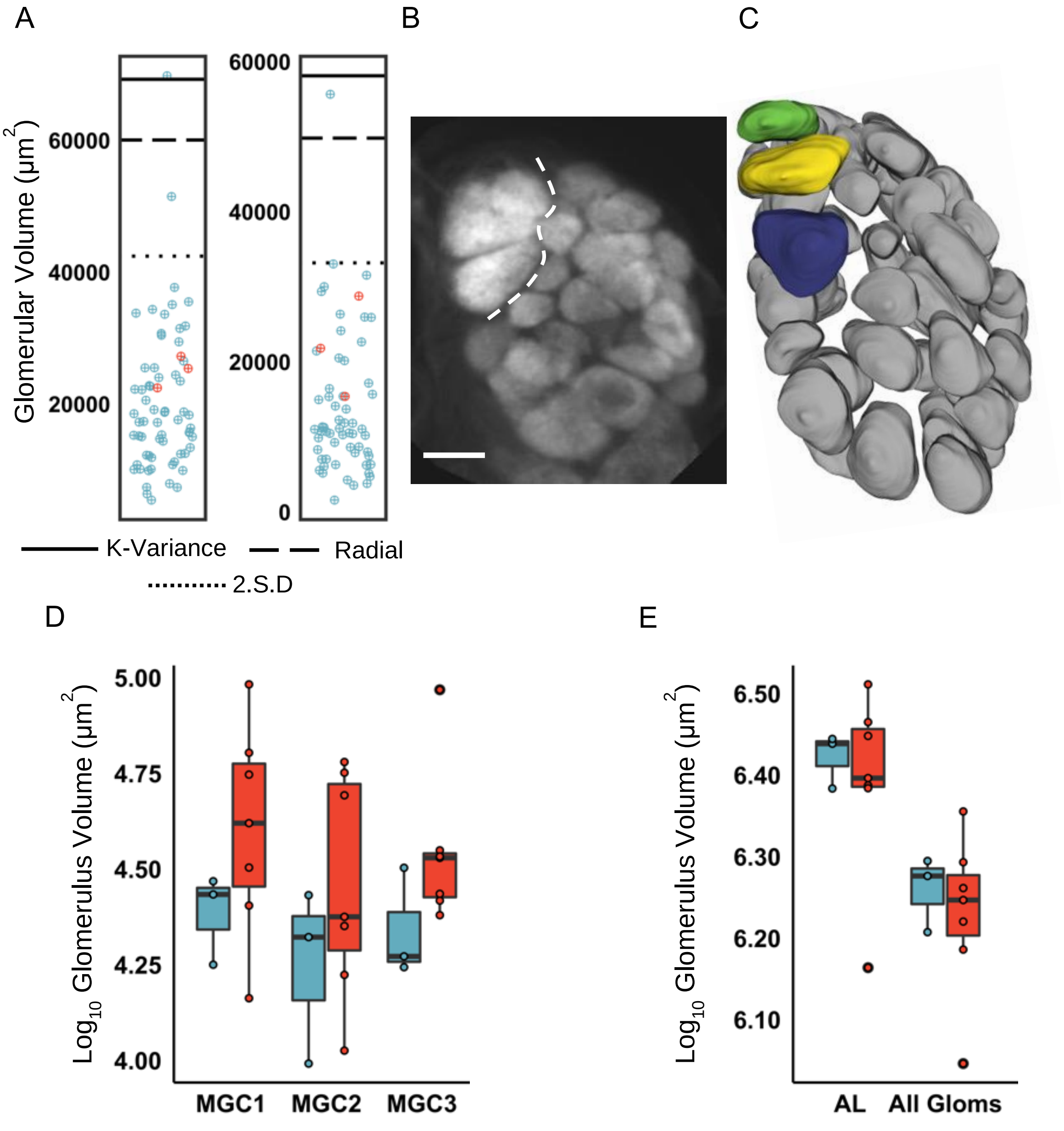


**Figure S9:** ***Oleria*** **MGC (♀5, ♂7) A)** Scatterplot displaying individual glomerular volumes of two focal males. Glomeruli that are part of the putative MGC in Red, all other glomeruli Blue. Cut off lines indicated. **B)** Confocal image of *Oleria* AL. MGC are deliminated from the rest of the AL by dashed line. Scale bar = 50µm. **C)** Surface model showing segmented glomeruli from a focal male. MGC glomeruli coloured, other glomeruli in grey. **D)** Boxplot displaying volumes of female (blue) and male (red) MGC components. **E)** Boxplot showing volumes of AL and total non-MGC glomerular volumes of female (blue) and male (red). Significance from linear models shown by: *<0.05, **<0.01, ***<0.001.

#### *ix) Hyposcada*:

We sampled five female and seven male *Hyposcada illnissa* individuals in this study. We do not observe any MG in this genera (Figure S10 A). We identify two glomeruli that appear separate from other AL glomeruli both though position and internal appearance (Figure S10 B, C). Whilst we consider these glomeruli as analogous to the ithomiine MGC for our analyses, they more strongly resemble the putative pheromonal glomeruli of *D. plexippus* (Heinze and Reppert, 2012). Neither of these glomeruli are seen to be sexually dimorphic.

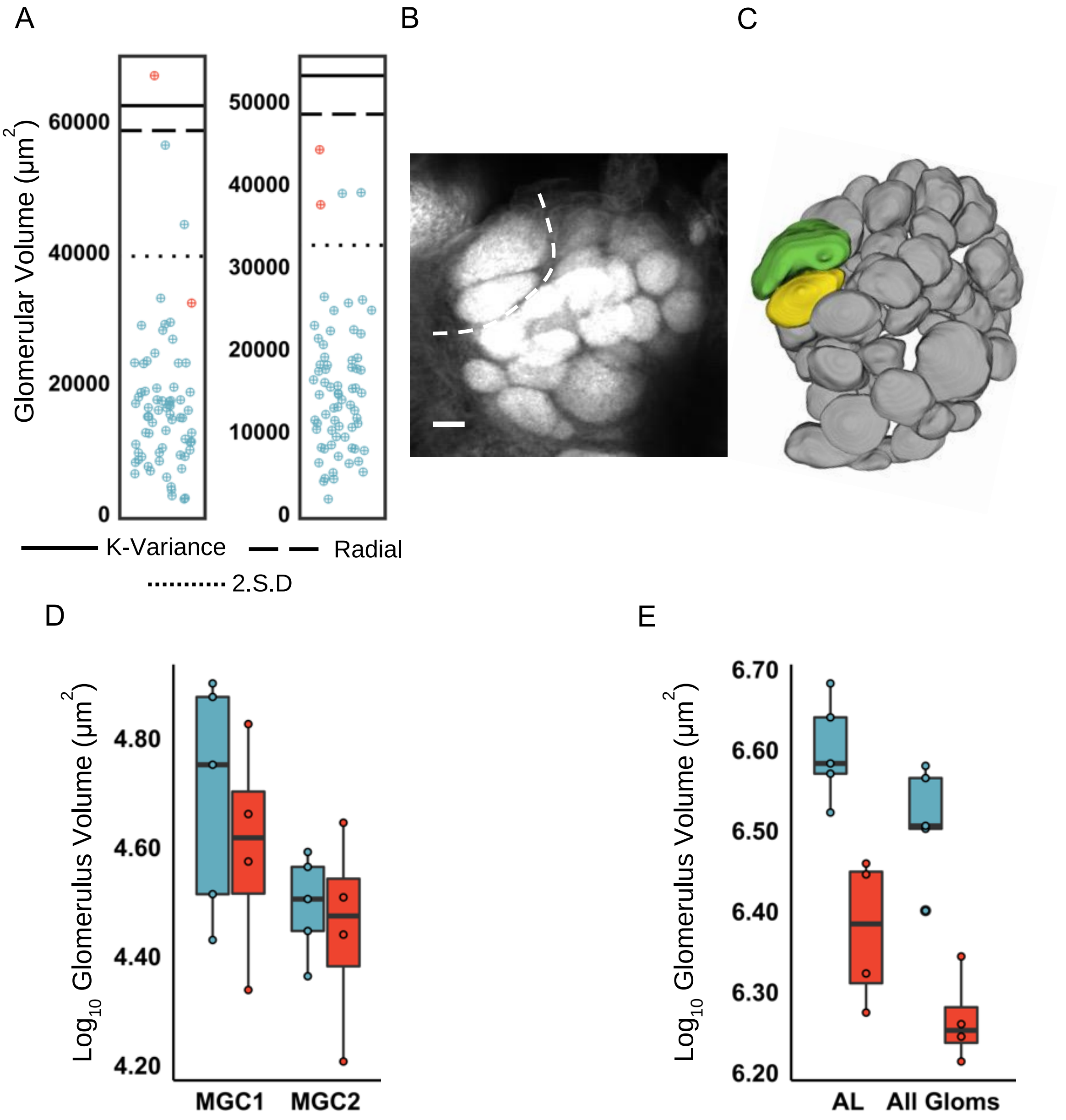


**Figure S10: Hyposcada MGC (♀5, ♂7) A)** Scatterplot displaying individual glomerular volumes of two focal males. Glomeruli that are part of the putative MGC in Red, all other glomeruli Blue. Cut off lines indicated. **B)** Confocal image of *Hyposcada* AL. MGC are deliminated from the rest of the AL by a dashed line. Scale bar = 50µm. **C)** Surface model showing segmented glomeruli from a focal male. MGC glomeruli coloured, other glomeruli in grey. **D)** Boxplot displaying volumes of female (blue) and male (red) MGC components. **E)** Boxplot showing volumes of AL and total non-MGC glomerular volumes of female (blue) and male (red). Significance from linear models shown by: *<0.05, **<0.01, ***<0.001.

#### *x) Callithomia*:

We assessed six male and five female *Callithomia lenea*. No MGs are identified in this genus (Figure S11 A). However, we observe a distinct glomerular complex, composed of four glomeruli (Figure S11 B, C). None of these glomeruli are sexually dimorphic (Figure S11 D, E).

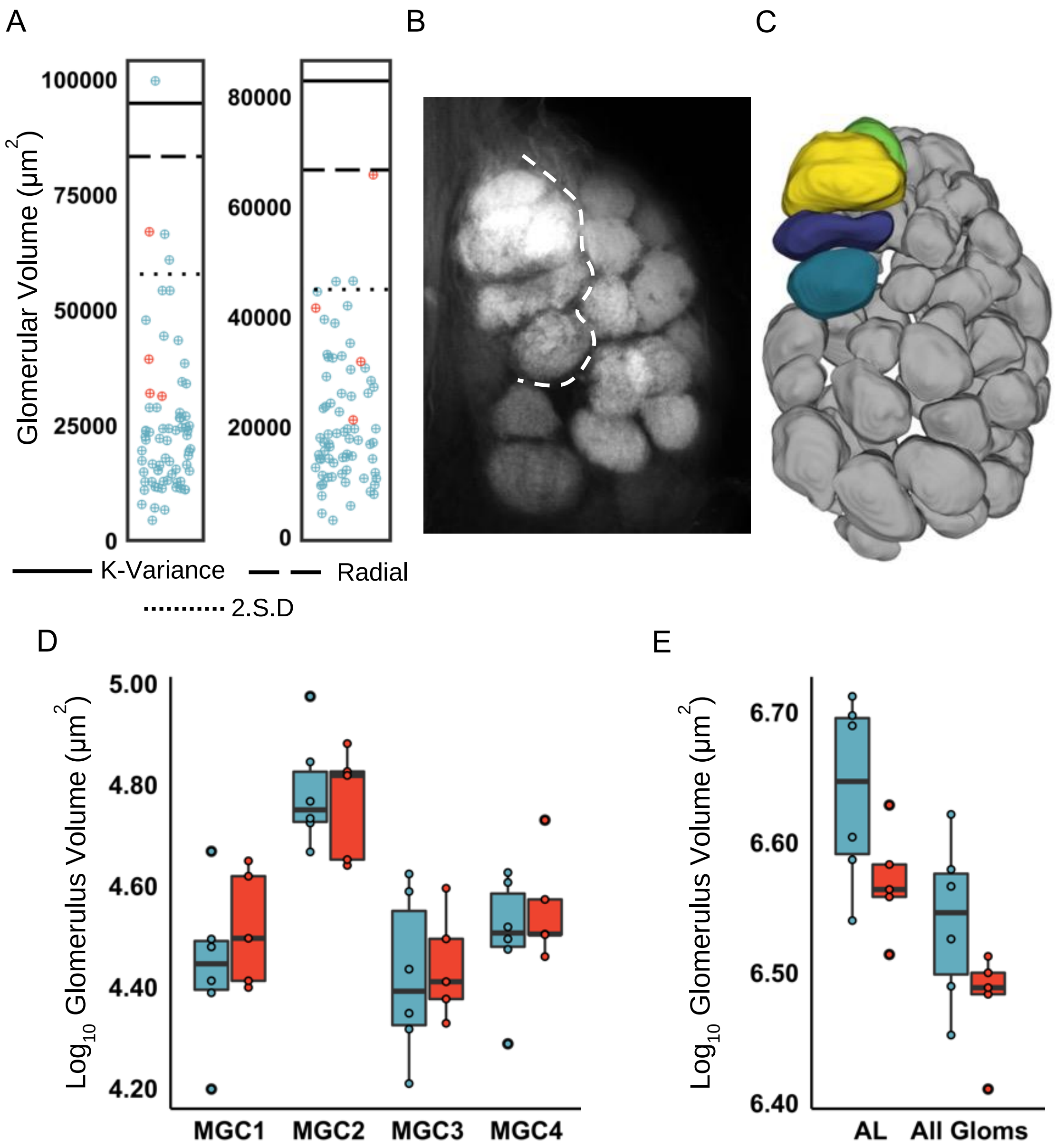


**Figure S11: Callithomia MGC (♀5, ♂6) A)** Scatterplot displaying individual glomerular volumes of two focal males. Glomeruli that are part of the putative MGC in Red, all other glomeruli Blue. Cut off lines indicated. **B)** Confocal image of *Callithomia* AL. MGC are deliminated from the rest of the AL by a dashed line. Scale bar = 50µm. **C)** Surface model showing segmented glomeruli from a focal male. MGC glomeruli coloured, other glomeruli in grey. **D)** Boxplot displaying volumes of female (blue) and male (red) MGC components. **E)** Boxplot showing volumes of AL and total non-MGC glomerular volumes of female (blue) and male (red). Significance from linear models shown by: *<0.05, **<0.01, ***<0.001.

### *xi) Pseudoscada*:

Eighteen *Pseudoscada florula* samples were evaluated, seven males and eleven females. Two MGs are identified in this genus (Figure S12 A). Whilst one of these MGs is part of a distinctive MGC (MGC1), the second is not. This is the only case among sampled ithomiines where we have identified a MG outside of the MGC. The MGC is particularly pronounced in this genera, and is composed of a aforementioned MG alongside four satellite glomeruli (Figure S12 B, C). We observe a large degree of sexual dimorphism in the MGC, with the MG and two satellites sexually dimorphic (MGC1, MGC2, MGC3) ( Figure S12 D, E).

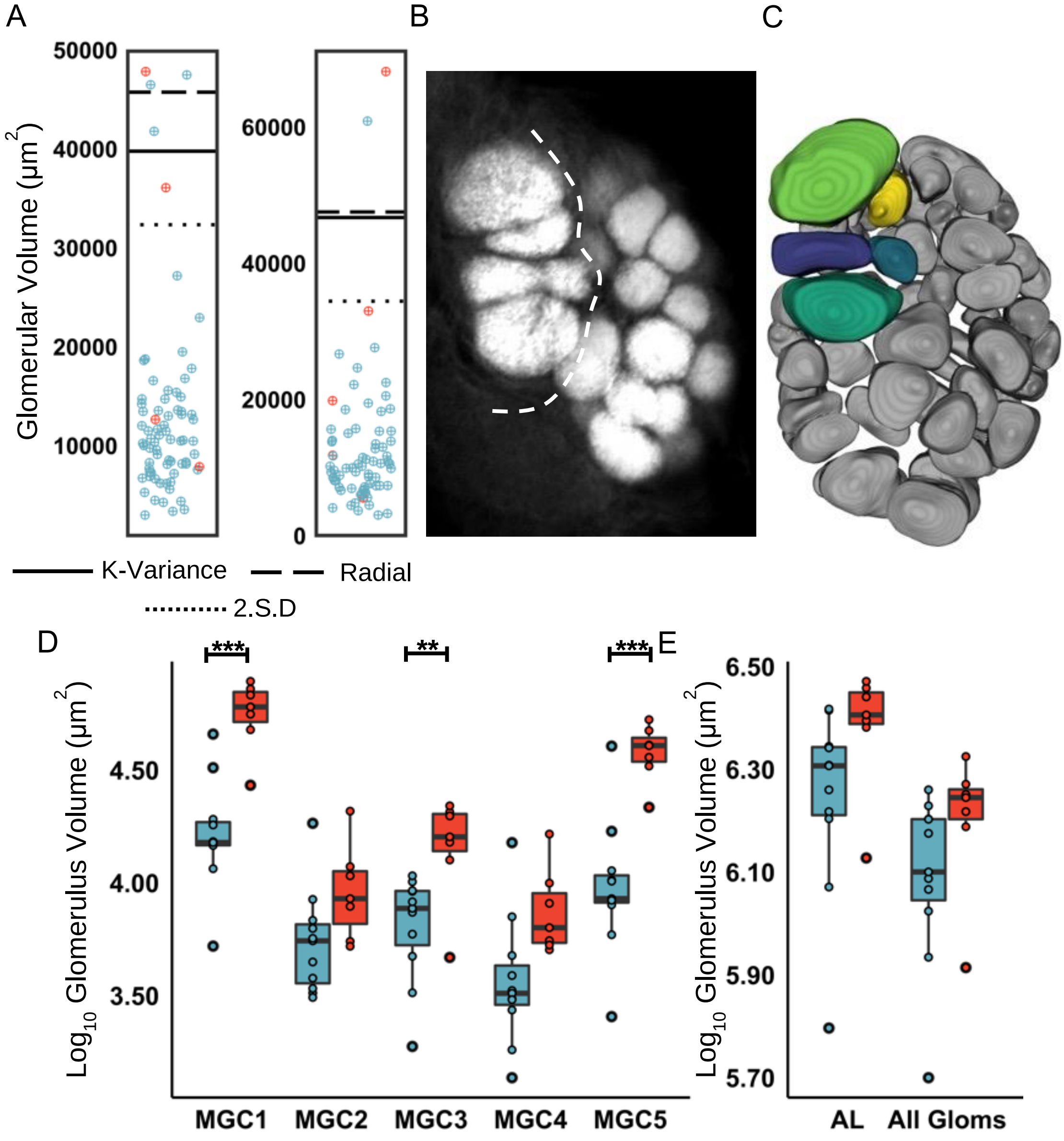


**Figure S12: Pseudoscada MGC (♀11, ♂7) A)** Scatterplot displaying individual glomerular volumes of two focal males. Glomeruli that are part of the putative MGC in Red, all other glomeruli Blue. Cut off lines indicated. **B)** Confocal image of *Pseudoscada* AL. MGC are deliminated from the rest of the AL by a dashed line. Scale bar = 50µm. **C)** Surface model showing segmented glomeruli from a focal male. MGC glomeruli coloured, other glomeruli in grey. **D)** Boxplot displaying volumes of female (blue) and male (red) MGC components. **E)** Boxplot showing volumes of AL and total non-MGC glomerular volumes of female (blue) and male (red). Significance from linear models shown by: *<0.05, **<0.01, ***<0.001.

### ***xii) Hypoleria***:

A total of sixteen *Hypoleria sarepta* samples were assessed in this work, six females and ten males. A single MG is identified in this genus (MGC2) (Figure S13 A). This forms a MGC in addition to three satellite glomeruli (B, C). We note that there appears to be some substructures apparent in MGC2, however they we're not clearly identifiable as distinct structures (Figure S13 B). We do not identify any sexual dimorphism in the MGCs constituent glomeruli, however MGC4 had a moderate effect size (Hedges *d*: MGC4 = 0.870) ( Figure S13 D, E).

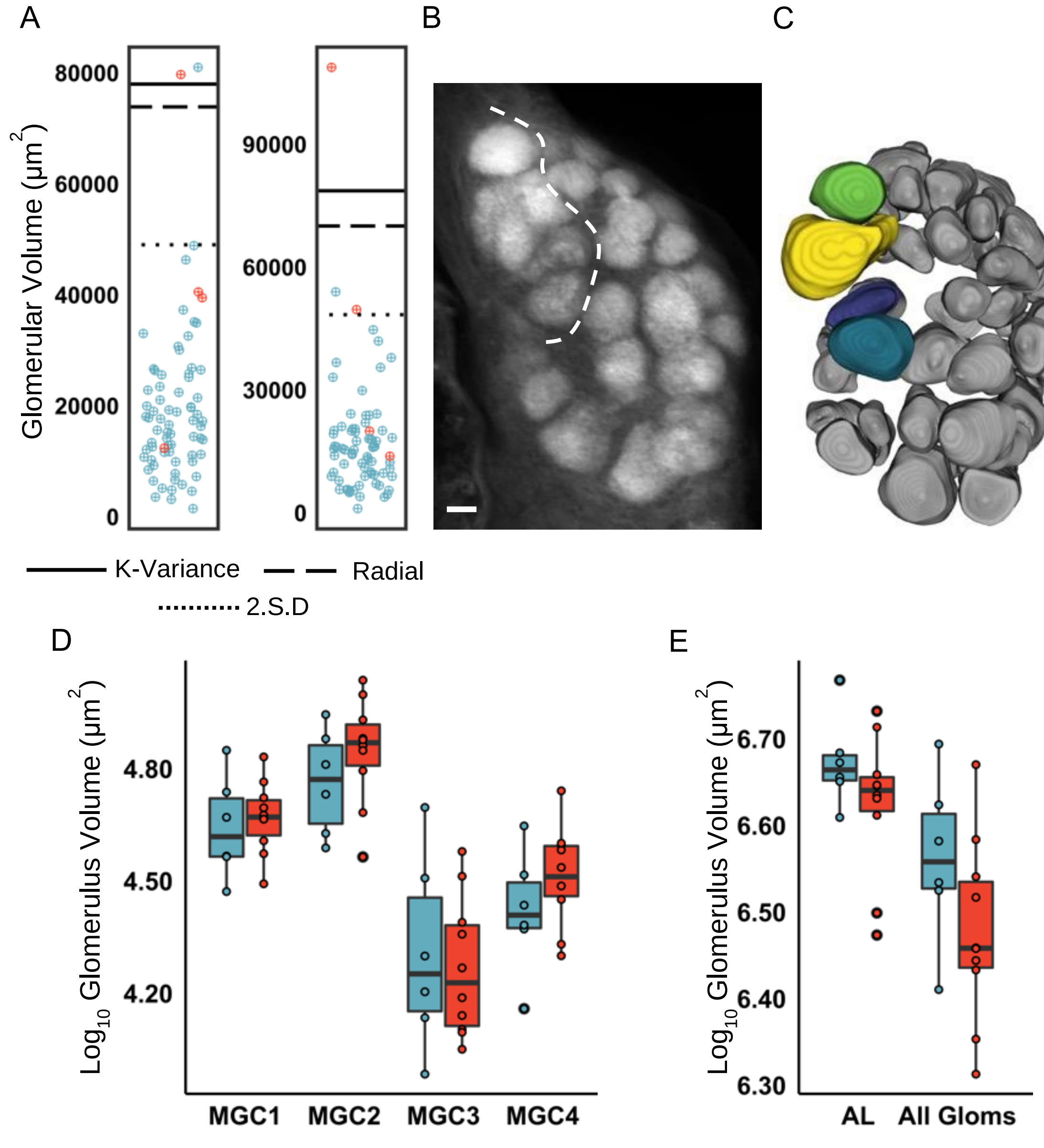


**Figure S13:** ***Hypoleira*** **MGC (♀6, ♂10) A)** Scatterplot displaying individual glomerular volumes of two focal males. Glomeruli that are part of the putative MGC in Red, all other glomeruli Blue. Cut off lines indicated. **B)** Confocal image of *Hypoleria* AL. MGC are deliminated from the rest of the AL by a dashed line. Scale bar = 50μm. **C)** Surface model showing segmented glomeruli from a focal male. MGC glomeruli coloured, other glomeruli in grey. **D)** Boxplot displaying volumes of female (blue) and male (red) MGC components. **E)** Boxplot showing volumes of AL and total non-MGC glomerular volumes of female (blue) and male (red). Significance from linear models shown by: *<0.05, **<0.01, ***<0.001.

#### xiii) *Godryis*:

We reevaluated the *Godyris zavaleta* data of Montgomery and Ott (2015) which assessed eight males and eight females. Though a single MG has previously been reported (Montgomery and Ott, 2015), this glomerulus does not consistently reach the threshold used in this study for classification as an MG (Figure S14 A). Despite this, there is a clear glomerular complex that is raised and distinct to that of the rest of the AL (Figure S14 B, C). Within this, there are two sexually dimorphic components, MGC1 and MGC4 (Figure S14 D, E).

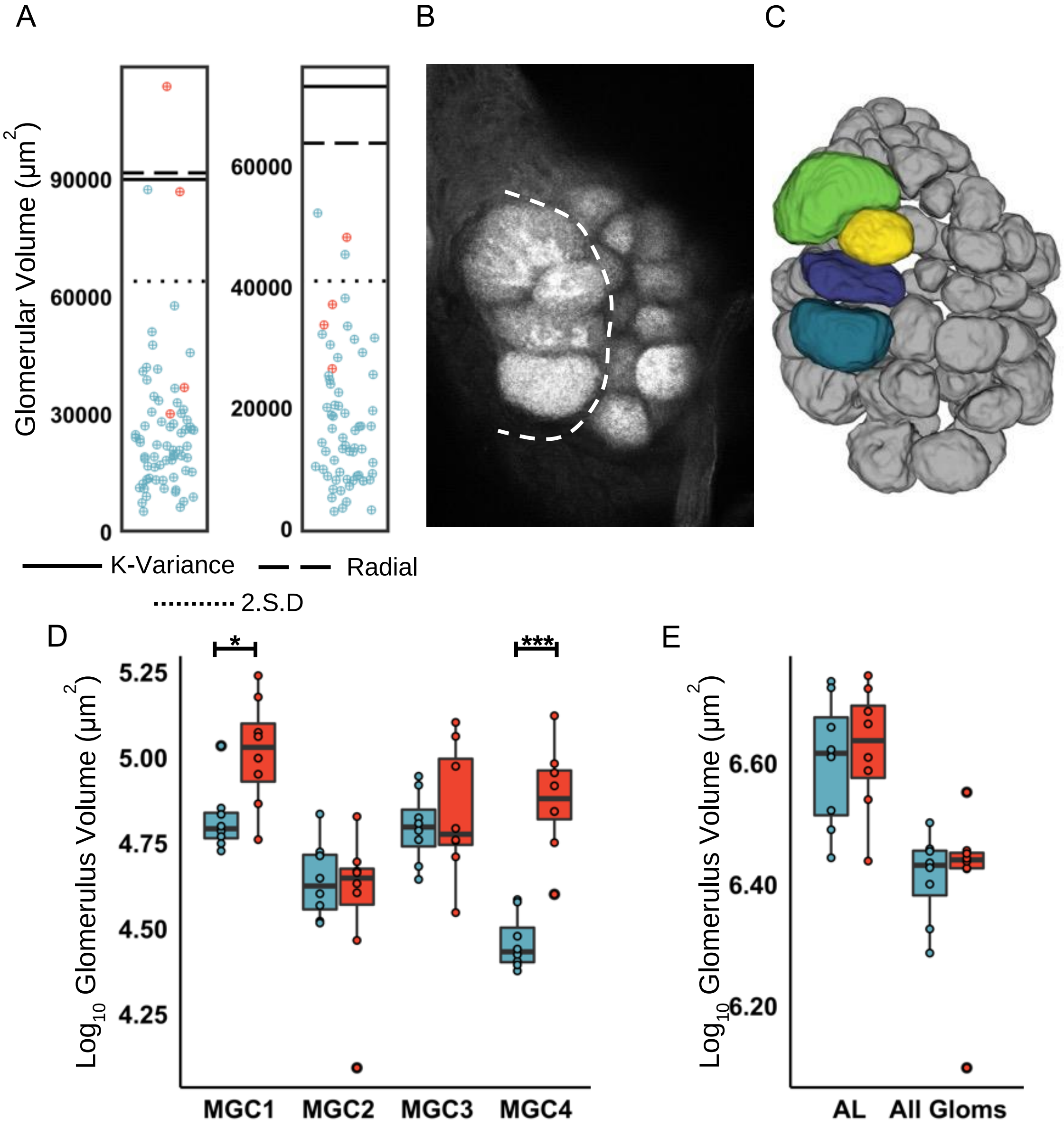


**Figure S14: *Godryis* MGC (♀8, ♂8) A)** Scatterplot displaying individual glomerular volumes of two focal males. Glomeruli that are part of the putative MGC in Red, all other glomeruli Blue. Cut off lines indicated. **B)** Confocal image of *Godryis* AL. MGC are deliminated from the rest of the AL by dashed line. Scale bar = 50µm. **C)** Surface model showing segmented glomeruli from a focal male. MGC glomeruli coloured, other glomeruli in grey. **D)** Boxplot displaying volumes of female (blue) and male (red) MGC components. **E)** Boxplot showing volumes of AL and total non-MGC glomerular volumes of female (blue) and male (red). Significance from linear models shown by: *<0.05, **<0.01, ***<0.001.
